# Inference and validation of causal genetic interactions in aging human fibroblasts

**DOI:** 10.64898/2026.10.21.760791

**Authors:** Madison S Krieger, Denitsa Milanova, David Thompson, Nadire R Cochran, Tongyao Wang, Barsha Pokharel, Robert P Ford, Andrei H. Gheorghe, William Gilpin, Yoav Freund, George Sugihara, George Church

## Abstract

In the past hundred years, the fraction of fatalities due to age-related disease has risen from an envied minority of cases to the leading cause of death and economic morbidity in the developed world. Despite notable advances in understanding aging, its intrinsic complexity has prevented major therapeutic breakthroughs. This complexity extends across all systems and scales in an organism, leaving a ‘fingerprint of aging’ across every aspect of cell and tissue behavior whose interrelationships are difficult to untangle. Compounding this problem, human aging is observed largely through cross-sectional rather than longitudinal data. Whether such data retain sufficient dynamical information to reveal causal interactions is unclear. Here we show that they can. Using convergent cross mapping (CCM), a method for identifying causal interactions in nonlinear dynamical systems, we reconstruct directed genetic interactions from age-ordered, cross-sectional transcriptomic data—recovering dynamics from populations rather than individuals. Applied to dermal fibroblasts from 490 GTEx donors, CCM identifies a genome-scale causal network whose topology, together with age-dependent changes in gene expression, predicts candidate drivers of aging. We tested these predictions experimentally by single-gene overexpression in fibroblasts from 55-year-old donors. Single-cell RNA sequencing confirmed 53.8% of testable CCM-predicted causal interactions, showing that dynamics reconstructed from cross-sectional populations contain experimentally recoverable information about gene-gene interactions. This 53.8% additionally exceeded that produced by multiple comparanda methods and a random null. We then asked whether this information could identify interventions that reverse cellular aging. Independent transcriptomic and high-dimensional morphological assays show that CCM-selected perturbations shift old fibroblasts toward states characteristic of young cells. These two measures of rejuvenation are strongly correlated, and the strongest interventions cause a majority of 55-year-old cells to resemble cells from 25-year-old donors. Thus, causal dynamics can be recovered from cross-sectional population data and used to identify interventions in aging. More generally, the results suggest a way to distinguish genes that change with age from genes that drive those changes, providing experimentally testable targets for altering the aging phenotype.

## Introduction

The social and economic impact of aging is under ever-increasing scrutiny, as both life expectancy and average age increase for many populations in the developed world. The average American citizen will invoke nearly $60,000 of medical bills in their last year of life, of which roughly 65% will be covered by programs such as Medicare.^1^ A recent estimate has placed the economic value of increasing the average lifespan by even 1 year at a staggering $38 trillion dollars.^2^

Concomitant with this rise in cost has been increasing scientific interest in aging as a multi-system, multi-scale disease state, implicating functions ranging from sub-cellular organelles to entire tissues. While many critical discoveries have concerned mechanisms of aging at the level of the entire genome, such as the role of telomerase^3^ and histone methylation,^4^ focus has recently turned to the importance of particular genes, gene networks, and pathways in controlling both longevity (lifespan) and the progression of aging phenotypes.^5–7^ Many of the difficulties in discovering key genes, pathways, and interactions with specificity to aging are obvious. For instance, the sheer number of genetic elements^8^ and regulatory mechanisms^9^ immediately reveals the size of the combinatorial search space in which to locate the keys to understanding aging biology. Additionally, there is growing understanding of not only the breadth of inter-human variability in the aging process, but even intra-human; many single-cell studies which attempt to quantify aging have found large variability – even amongst cells harvested from the same human donor.^10^ Finally, there is the challenge of collecting empirical data on human aging. Despite admirable attempts,^11^ there is an extreme lack of longitudinal aging studies at the level of detail and richness expected for disease-specific multi-omics studies.

We believe that between-donor variability and sparse measurements may present an opportunity, rather than purely an obstacle. Because age is one of the must abundantly available pieces of *meta*data in large-scale repositories such as GTEx,^12^ access to troves of multi-omics data covering all ages in a normal human lifespan is easily obtainable. In some senses, any dynamic information on aging which can be derived from these sources could even be advantaged in comparison to longitudinal data. For instance, the idea that aging can proceed along several distinct dynamical trajectories “ageotypes”^11^ suggests that diverse, large-scale repositories may contain a broader perspective than would be obtainable within a single individual’s longitudinal ageotype.

Here we propose that the combinatorial problem of inferring gene-gene interaction networks with specificity to aging, as well as determining the amount of usable signal in non-longitudinal aging data, can be probed using the same set of tools. We draw our tools from the growing *empirical dynamics* literature – applying principles of nonlinear dynamics to empirical time series data, but in a way that does not require explicit modeling assumptions on the processes that generated the data. We view the challenge of finding actionable interventions in aging as a Tukey paradigm^13^ – the principal goal is an informed search through an impossibly large combinatorial space (“explore”), combined with experimental feedback on the search procedure (“confirm”) and, eventually, deeper profiling on the biological “why” of positive hits emerging from the search (“explain”).

Our primary search tool is convergent cross-mapping (CCM),^14^ which aims to provide a binary yes/no response to the question “Does the time series of variable X quantitatively affect the time series of variable Y?”. Here, the variables are gene expression, and the time series is cobbled together from non-longitudinal data, a process which we call *quasi-time series*. When applied to a single pair of genes X and Y, the validity of both the method and the underlying data composition techniques can be examined to an extent experimentally (for instance, by overexpressing X and observing if Y is differentially expressed as a consequence). At a larger scale, when applied to all distinct pairs of genes in a dataset, we argue that more impactful inferences can be made about which genes are *most likely* to produce some sort of rejuvenative effect if perturbed. This is a much harder problem, and one which we believe to be much less explored. We therefore try to adopt the simplest approach to identifying fruitful targets, instead focusing our efforts on rigorously quantifying the performance of these targets; enhancing and optimizing the actual target discovery problem is largely left as an open question.

As a pilot investigation into these questions, we applied CCM to *quasi-* time series generated from a large dataset (GTEx v10) of dermal fibroblast bulk RNA measurements. Dermal fibroblasts are appealing for at least two reasons – first, they are one of the most accessible cells in terms of public data (for instance, GTEx curates only one other isolated cell line) as well as being available from diverse donors in many biobanks and representing a lower challenge to culture and probe experimentally. Secondly, fibroblasts are generally considered to be a reasonable paradigm for aging,^15^ exhibiting many of the broader “hallmarks of aging”^16^ that are typically cited as pan-tissue effects taking place during the aging process. This enhances our ability to connect our results to the broader aging literature.

In what follows, we describe how we generate our *quasi-* time series, as well as how we apply CCM to generate a very large network of imputed interactions between genes during the process of aging. We use an extremely simple heuristic on this network to rank targets in terms of their putative rejuvenative potential. We then experimentally probed both the per-gene-relationship accuracy of the CCM + *quasi-* time series combination, as well as broader questions about the rejuvenative impact of gene expression perturbations, using library screens in scRNA and single-cell microscopy applied to mixed cells representing an older (55-year-old) and younger (25-year-old) population. We suggest a novel machine-learning based method for quantifying rejuvenation, which experimentally recapitulates many hallmarks of aging. The per-gene-interaction results strongly suggest that there is value in abundantly-available *quasi-* time series, and that tools like CCM are a reasonable starting point to extract this information. The results of our library screens suggest that, while there is obvious room for improvement in how one determines rejuvenative targets from this wealth of information, even our simple heuristic will reliably find targets of interest; according to most of our experimental metrics, our best-performing library member causes a majority of transduced cells to be more akin to 25-year-old cells than to 55-year-old cells.

### Creating human transcriptomic *quasi-* time series from public data

The empirical data used to infer gene-gene interaction networks was taken from the “Cells – Cultured fibroblasts” subset of GTEx^17^ (V10), consisting of dermal fibroblasts from 490 donors with ages ranging from 20 to 70 (Fig. 1**(a)**). After transcripts-per-million (TPM) normalization and quality control procedures, the library consisted of 5,991 measured gene expression levels.

**Figure 1.**
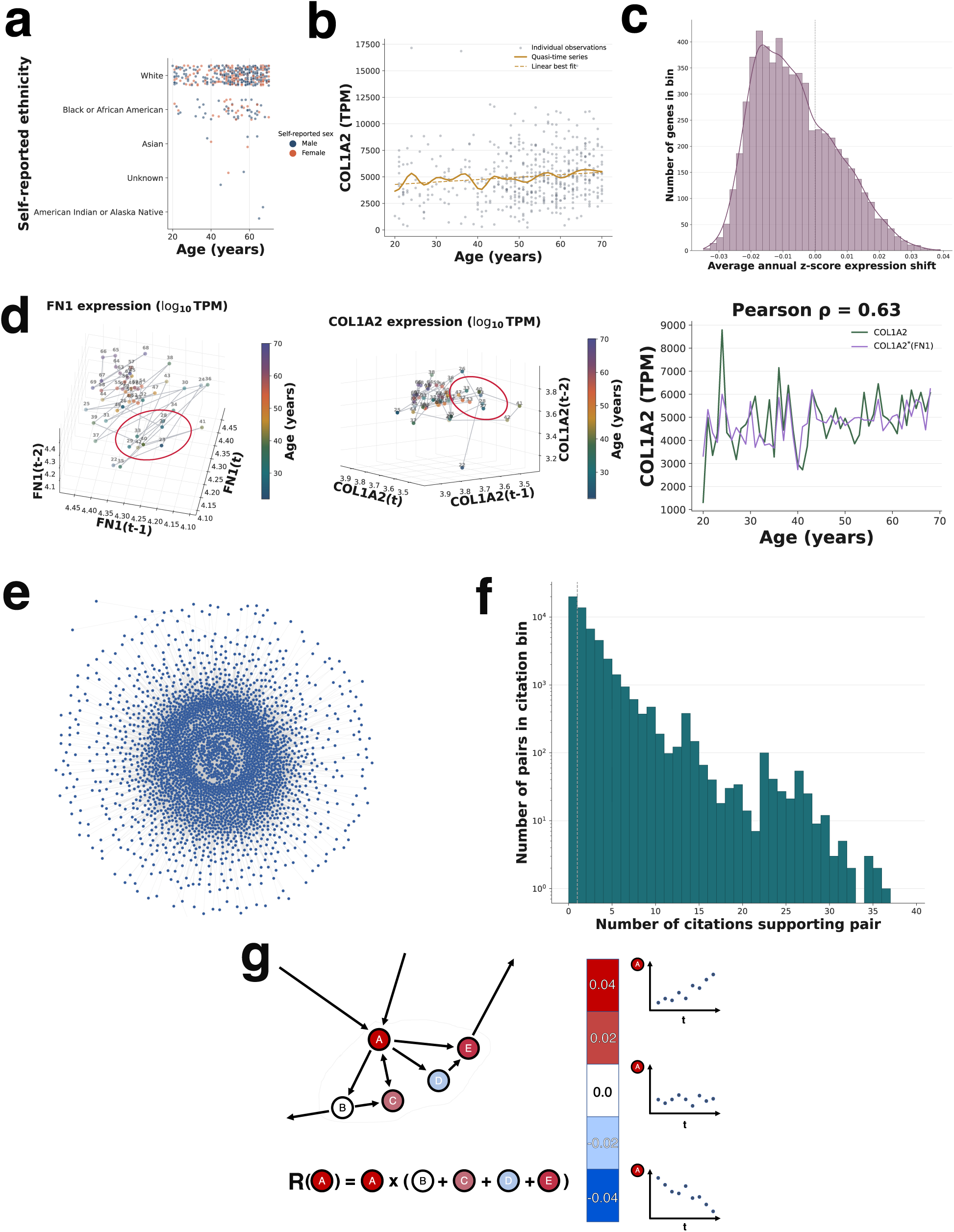
Inferring mechanistic interactions in the transcriptome. **(a):** Population demographics within the GTEx “Cells - Cultured fibroblasts” tissue domain. **(b):** For each gene, we form a single “quasi” time series consisting of every integer year of life from 18 to 75, and a single expression value per year, taking the average over the expression for all donors of that age. Here, a Gaussian filter with *σ* = 1.5 was used for smoothing, for visualization purposes only. **(c):** Taking the *z*-score of each genic quasi-time-series and fitting a line indicates the general trend of that gene’s expression with age; in fibroblasts, a majority of genes decline in expression by approximately 1 *z* over the human lifespan. **(d):** CCM^14^ infers a connection between two time series if local neighborhoods of *delay embeddings* are correlated. Here we show a three-dimensional delay embedding of the time series representing FN1 (fibronectin) and COL1A2, which we find to be interacting due to the correlation of time-indexed points which spatially cluster in each gene’s delay embedding. This is a known interaction – depriving the ECM of fibronectin deactivates the integrin-mediated YAP/TAZ pathway, which leads to decreased secretion of collagen.^29,30^ Red ovals indicate examples of ages that cluster together in delay embeddings. When this clustering is robust, prediction of one gene’s expression (here, COL1A2) based on clustered neighbors of another gene expression (FN1) is highly accurate and an interaction is assumed. **(e):** Repeating this process for every possible gene-gene pair yields a set of directed interactions, which, when pictured as arrows from the influencing gene to the influenced, forms a large directed network. **(f):** A histogram indicating, for each gene-gene pair in **(e)**, how many supporting citations were found in the BioGRID resource. More than half of the directed edges have at least one supporting citation. **(g):** Our *ad hoc* initial heuristic for ranking the relative importance of genes in driving the aging process in dermal fibroblasts. The ranking score for a putative gene *A* is determined by an intrinsic component (the linear fit slope of the *z*-transformed quasi-time-series of *A* described in **(c)**), times an extrinsic component formed by the sum of the same slopes for every gene receiving a directed arrow from *A*.

We then construct“*quasi-* time series” for each gene. We bin donors by their integer-valued age (between 20 and 70) and average the TPM value of each gene across all donors within a given age bin. Our approach follows prior approaches to extracting ground truth temporal information from cross-sectional studies .^18^ An example of a *quasi-* time series, that for COL1A2, is shown in Fig. 1**(b)**.

Note that there is a strong bias in *quasi-* time series towards a decline in expression with age (Fig. 1**(c)**). This comports with other studies on aging fibroblasts, which suggest that broad classes of functional genes (such as those encoding extracellular matrix proteins) decline in expression as a function of age.^19,20^

### CCM unravels a vast network of literature-supported transcriptomic interactions

We suspect that the relatively low degree of investigation into how simple objects like our *quasi-* time series can be used to probe the time dynamics of aging is due to the incredibly noisy nature of individual-gene signals. Traces such as that in Fig. 1**(b)** indicate highly nonlinear behavior, in which local changes in expression arise from known within-cohort variance.^21^ This variance itself is known to increase during the aging process.^22,23^ We argue instead that there is usable information in *quasi-* time series that reflects population-aggregate aging dynamics, which can be extracted using tools geared towards nonlinear dynamics and networks of causal interactions. We thus seek to infer mechanistic interactions between genes with convergent cross mapping (CCM),^14^ a model-free technique for inferring causal interactions from time series. CCM has previously been applied to sparse and noisy time-series data arising in ecological and physical systems,^24,25^ and in small proteomic networks has been effective in revealing experimentally-actionable interactions in diseases known for nonlinear proteomic behavior.^26^ We believe our reasoning is supported by recent advances in similar methods that attempt to extend the theoretical underpinnings of CCM to cross-sectional data.^27^ The method consists of a leave-one-out cross-validation test for Takens’ theorem,^28^ a result lying at the intersection of nonlinear dynamics and differential topology, which uses pairs of time series *X, Y* to create a prediction time series *X*^^^, predicated on the concept that *X* and *Y* are coupled via a deterministic dynamical system. If *X*^ converges to *X* as more and more data is used, then *Y* has a mechanistic or causal impact on *X*.

An example of a single calculation on two of our *quasi-* time series is shown in Fig. 1**(d)**. These two genes, FN1 and COL1A2, represent a known interaction in dermal fibroblasts – the stiffness of proto-ECM is determined by fibronectin density, which signals cells to begin synthesizing larger collagen fibrils via the integrin-mediated YAP/TAZ mechanotransductive pathway.^29,30^ Even though their time traces are very noisy, both when viewed in 2D and in higher dimensions (achieved by using lags of the *quasi* time series as different variables), the prediction COL1A2*\** generated from knowledge of FN1 and the assumption that the two interact is quite accurate (Pearson *ρ* = 0.63).

Applying this method at genome-wide scale allows us to infer a large network of gene-gene interactions (see **Methods**), producing in a network of 5,962 genes interacting via nearly 1.8 million directed edges (Fig. 1**(e)**). To validate this procedure, we first use BioGRID, a large repository of curated evidence for over 1.34 million distinct gene-gene interactions in humans.^31^ We query each of the 1.8 million CCM-inferred interactions against BioGRID, and record how many distinct pieces of literature in the database support the existence of the interaction in *H. sapiens* (Fig. 1f). We find that 54.9% of inferred interactions match at least one literature reference, and 14.2% of edges matched at least 5 references. This shows an enrichment in literature matches against a null distribution of the same size, drawn from all possible edges that might have been inferred; the null has 44.2% of inferred interactions matching at least one reference, and 6.3% matching five or more references (Fig. S2).

### Identifying genetic drivers of aging

While we are interested in the per-edge accuracy of our CCM-derived gene-gene interaction network as a base validation step for both the construction of *quasi-* time series and the application of CCM to this type of data, we believe a second and more powerful form of validation might come from analyzing and situating individual genes in the *global* network.

A gene’s *quasi-* time series, as well as its position in the network, render a substantial amount of information which we can use to rank genes in order of an imputed “importance” in the aging process. As a first step towards exploring how our methodology can render actionable data for the study of aging and rejuvenation, we develop the simplest possible ranking heuristic that incorporates as much of each gene’s available features, while also minimizing and inter-individual variation. Our chosen heuristic, the “RankScore”, consolidates the dynamic features of individual genes with the broader interaction network. The dynamic (or, “intrinsic”) featurization of gene *g* is given by *I_g_*= *S_g_*(1 *− p_g,Spearman_*), where *S_g_*is the slope of the best-fit line to *g*’s *quasi-* time series, and *p_g,Spearman_*is the *p*-value resulting from the Spearman correlation between the ages and expression values in the *quasi-* time series – a measure of the monotonicity. The absolute value of *I_g_* is small when a gene either changes very little with age, or has a particularly noisy trace. The network-derived (or, “extrinsic” featurization of *g* is simply the weighted outdegree – the sum of the above intrinsic features *I_j_* for all genes *j* receiving an edge emanating from *g*. The overall RankScore *R_g_*is thus given by

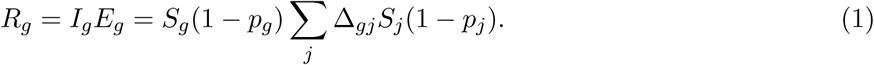

This heuristic balances local properties of each gene’s dynamics with its broader role within the network. Moreover, because the network featurization is limited to a gene’s immediate neighborhood (i.e., 1-step walks emanating from the focal gene), the RankScore is interpretable: genes with the highest absolute RankScore must generally show strong, reliable changes in their own expression level with age; they must furthermore be imputed by CCM to be driving the expression level of many other genes. This heuristic is visualized conceptually in Fig. 1**g**.

### Individual causal interaction validation via perturbed transcriptomics

Validating the millions of causal interactions inferred via CCM – which, putatively, take place on the timescale of years in the course of normal aging – represents a significant challenge. The logistical realities involved in empirically studying aging generally prohibit us from investigating the role of single-gene perturbations at the timescale of years *in vitro*, let alone *in vivo*.

However, we reason that any causal interactions between genes that are relevant to the aging process can be coarsely binned into two categories: interactions that also exist at faster timescales, and interactions that do not. From a modeling perspective, the former have Jacobian coefficients that are *time(scale)-independent* ; from a biological perspective, we might say these interactions are *intrinsic* or *necessary*. In other words, these interactions are <u>not</u> specific to the aging process, but constitute “normal” biology, and should therefore be capable of validation through normal assays. We then accept that any interactions which are *not* validated by normal assays exist in unknowable proportions of false positives – failures of the methodology – and true positives which obey a biology separate from the timescale of days or weeks, and which are therefore too difficult to validate given present experimental methods.

Because our underlying data is transcriptional, our main validation assay is single-cell RNA readouts from dermal fibroblasts which have been perturbed via single-gene overexpression. The primary goal is to evaluate the validity of directional causal interactions, where directed edges in the causal network stem from the overexpressed library member and terminate on genes whose expression level is measured by the assay. However, since the ultimate goal of the computational approach is to evaluate targets in the context of aging and rejuvenation, higher-order outputs, such as inferred phenotypic changes (which are strongly implicated in the aging literature) are also deeply important.

### Designing a lentivirus overexpression library for validation experiments

We perturb the transcriptome of individual cells via overexpression, induced through additional plasmid copies of single-gene library members delivered with a lentiviral vector. Overexpression – as opposed to silencing RNA or editing-driven knockout – is motivated by the underlying biology of fibroblasts. Skin, like many tissues, generally shows a decline in transcriptomic activity,^19,20^ with the majority of genes expressed at lower TPM with increasing age (Fig. 1**(c)**). The most potent interventions might therefore seek to reverse this steady decline.

We design our library based on the CCM results, deferring further consideration based on known biological pathways (such as whether they represent transcription factors, secreted factors, and other properties that might motivate a different optimal experimental protocol). While there are a variety of outputs from the computational analysis – such as local or global network properties (degree distribution, centrality, and so forth) – we reasoned that we could best evaluate both the underlying method for identifying causal interactions, as well as the heuristics for ranking the rejuvenative potential of targets (Equation 1), by binning the library based on the latter ranking. We create three bins: genes with the highest overall RankScore, the lowest overall RankScore, and genes with RankScore very close to 0 (Fig. 2**(a)**).

**Figure 2.**
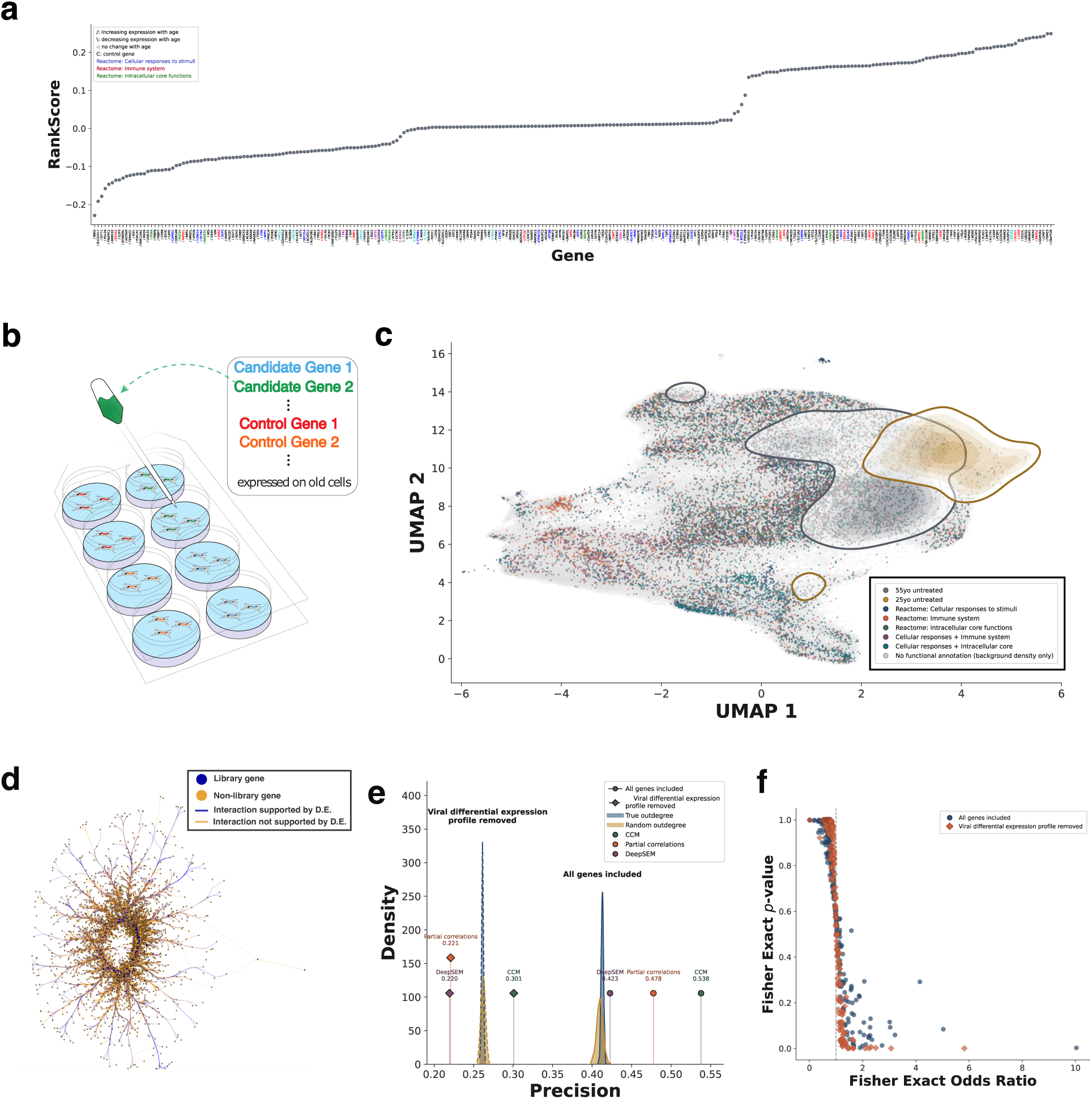
Single-cell RNA profiling of an EDM-inspired library. **(a):** Our ranking heuristic for a gene’s impact on the aging process (Fig. 1(g) assigns each gene a value: negative values typically imply strong driver genes (a large extrinsic factor) which are themselves increasing in expression with increasing age; high positive values typically indicate strong drivers whose expression decreases with age. We aimed to construct a genetic library consisting of roughly 100 genes from three bins of this ranking score: the lowest, the highest, and the ‘most neutral’ (closest to zero). Genes labelled on the *x*-axis have two characteristics besides their HUGO gene symbol: a text prefix, with “/” indicating that the gene increases in expression with increasing age, “′’ increasing the opposite, “C” indicating a control, and “-” indicating little-to-no change in expression with age. Primary colors indicate genes binned in (super)classes in the Reactome pathway enrichment system: red indicates genes along the immune axis, blue indicates cellular responses to simuli, and green indicates core cellular functions. Secondary colors indicate overlaps of the functional classes assigned primary colors. **(b):** scRNA profiling was performed on fibroblasts taken from arrayed wells, in which distinct wells received distinct transgenes from the library (including many untreated control wells). After QC on read quality, diversity, and other metrics, the number of distinct library members represented was 266. **(c):** A SAM^47^ projection of the 649,846 cells sequenced, using the same functional color scheme as **(a)**. **(d)** A subnetwork of the EDM network pictured in Fig. 1(e). This subnetwork consists of the 266 library genes (blue nodes), as well as all of the nodes in the full EDM network which: I) receive an arrow from the library nodes, and II) are indexed in the human genome build used for read alignment (see **Methods**). Blue edges indicate verification of the relationship via differential expression in the perturbed scRNA dataset; blue edges constitute 53.7% of all edges. **(e):** A precision comparison of the CCM methodology employed here to two alternatives – DeepSEM^52^ and partial correlations^50,51^ – as well as a permutation test representing the null hypothesis, wherein edges are drawn uniformly-at-random based on the CCM network degree distributions. In the hatched case, genes which are differentially expressed in the GFP-transduced cells (the “viral differential expression profile”) are removed from consideration. **(f):** The global network precision depicted in **(d)** is not equidistributed over all library genes. Here we report the odds ratio (test statistic) and associated *p*-value for a Fisher Exact test performed for each library gene, where the contingency tables consist of I) the observed genes differentially expressed when the library member is perturbed, and II) the predicted genes receiving an edge emanating from the library member in the EDM network. An odds ratio of 1 corresponds to the overlap between the two lists expected by random chance.

In addition, we include experimental controls (such as GFP, CRE and luciferase) as well as benchmarking controls, genes whose role in aging and rejuvenation is strongly implied in the current literature. These include NFE2L2 (NRF2),^32^ a transcription factor with known regulatory impact on cellular metabolism; SOD1,^33^ a gene whose deficiency causes senescence due to mitochondrial failure; TERT,^34^ telomerase reverse transcriptase; TP53, for its role in senescence; the metabolic factor AKT1;^35^ the cell adhesion factor ANXA2;^36^ and two negative controls: MeCP2,^37^ a silencing factor strongly associated with senescence, and DAP,^38^ a negative regulator of autophagy, a pathway whose dysfunction is thought to be a hallmark of aging. In addition, we annotate seven genes as controls which are not mentioned in the literature, but which were positively indicated during a similar pilot study in which we applied CCM to a smaller dataset:^39^ BGN, CLTA, CTNNB1, MAP4, PRRC2B, RPL19, and RPS23.

The logistical necessities of the experiment – such as optimizing and administering viral titer for each individual library member, to a distinct subpopulation of cultured fibroblasts – constrain the ultimate size of the library. We initially aimed to include roughly 100 genes in each of the three computationally-motivated bins, plus controls. After successive quality control procedures for virus manufacture, transduction (with drug selection for positive transfection), and scRNA sequencing, the final number of distinct library members present in the scRNA analyses is 266.

We believe it is noteworthy that the vast majority of genes in our library are not transcription factors – in fact, no large group of genes belongs to any functional group (such as transcription factors, secreted factors, and surface markers) associated with known signalling cascades in the field(s) of transcriptomic regulation or aging. To rigorously count these, we use large databases of human secreted factors,^40^ transcription factors,^41^ and surface markers.^42^ In resources that combined established markers with computationally-imputed ones, we exclude any markers that did not have experimental verification. Of the 244 non-control genes in the library, we have only 13 transcription factors ^1^, 13 surface markers ^2^, and 7 secreted factors ^3^. Note that in this annotation scheme, extracellular matrix (ECM) proteins are not considered as secreted factors. Our genetic library, with all meaningful annotations and many key results and summary statistics from following sections, can be found in the released codebase – see Appendix A.

### Generating a perturbed scRNA dataset for causal edge confirmation

Because of our focus on aging, we seek to balance a donor population of human dermal fibroblasts which could both resolve some aspect of the aging process as well as intra-cohort heterogeneity, without adding undue complexity and sources of noise due to differences in donor growth rate, response to culture medium, and other factors. We thus use a donor pool consisting of six donors, three aged 25-years-old (25yo) at time of donation, three aged 55-years-old (55yo) at time of donation.

These cells, save for a control population, are then arrayed into equally-sized wells, and the populations in each well are subjected to a single library member lentivirus overexpression perturbation (Fig. 2**(b)**; see **Methods** for experimental details). This procedure includes blasticidin-based drug selection for the transduced library member after transfection, in an attempt to ensure that only successfully transduced cells proceed to analysis.

After the transduction protocol and single-cell sequencing, a number of computational quality control steps are further applied. These generally follow standard procedures for removing cells from analysis based on count diversity or abundance, as well as ribosomal and mitochondrial count densities.^43^ Based on growing evidence that nuclear-bound lncRNA count densities represent a critical QC metric, we also included the key marker MALAT1 in our quality control measurements.^44–46^ While we do not explicitly reject cells based on count density, we also pay close attention to the roles of other nuclear lncRNAs, such as XIST and NEAT1, in our analytical readouts. The role of MALAT1 is treated analogously to mitochondrial reads: an aberrantly low count density likely indicates an empty droplet, lysed cell, or a cell with a heavily damaged nucleus; on the other hand, an aberrantly high count density likely indicates a damaged or otherwise weak plasma membrane, where increased count density probably indicates a loss of cytosolic RNA rather than a “true” abundance of MALAT1 transcripts within a healthy cell.

The resulting scRNA dataset consists of 649,846 cells distributed across 25yo and 55yo control populations as well as 55yo perturbed cells with perturbations ranging from 266 unique library members. A cursory examination of unsupervised embeddings (such as the SAM^47^ projection in Fig. 2**(c)**) fails to reveal obvious perturbation-driven clustering of treated cells, nor do we see extreme separation of control 25yo and 55yo populations. This indicates that the aging state is not directly apparent from expression levels alone, motivating the need for a computational analysis that considers dynamical state.

### Differential expression analysis validates half of CCM-inferred interactions

It is difficult to ascertain which inferred interactions might be considered “generic” (i.e., these interactions occur on non-aging-related timescales such as hours or days, and can therefore also be resolved on the timescale of years, albeit with an uninteresting relationship to aging), or which might be aging-specific. We reason that the scRNA screen should, with minor caveats (related to the *in vivo*, forced nature of our perturbations), replicate the “generic” interactions. A high number of replicated (validated) generic interactions would speak to the general usefulness and validity of the computational approach. For unvalidated interactions, it is unknown *a priori* which represent false positives, or true biology which fails to present itself in this assay either due to a mismatch in timescale (the interaction takes longer than weeks to manifest) or other unforeseen shortcomings of the assay.

We consider “validation” of a computationally-inferred gene-gene interaction to be the following: if we infer a directed edge from gene X to gene Y (meaning, the expression level of gene Y is affected or driven by the expression level of gene X), then we should observe differential expression of Y (compared to untreated 55yo control) in cells where gene X has been (massively) overexpressed via transduced plasmid. Here, we calculate differential expression between individual library member-transduced cells and untreated 55yo cells using a pseudobulked Welch *t*-test.^48,49^ We call a gene Y differentially expressed in cells transduced with X if the Bonferroni-adjusted *p*-value of the Welch *t*-test (with two-sided alternative hypothesis) is less than 0.05.

We find that the percentage of viable edges in the CCM network (the edges that were possible to validate; that is, directed edges emanating from a gene X belonging to the perturbation library and terminating on a gene Y both belonging to GrCH38 and existing in the original empirical time series data at measurable read counts) which are validated via differential expression was 53.8% (Fig. 2**(d)**). This percentage represents the number of inferred edges which also showed *X → Y* in differential expression analysis, divided by the total number of viable edges as defined above. This number is quite close to the 54.9% of interactions which were found in BioGRID, despite the difference in scope – the differential expression analysis is restricted to a small subset of the total CCM network, whereas BioGRID is queried for every single edge in the network. Interestingly, the overlap between these two categories is slightly lower than anticipated; only 62.1% of edges confirmed by differential expression are also present in the BioGRID query results.

In situating this result, it is critical to remember that the percentage of gene pairs which truly do interact (in generic biology, aging biology, or any other setting) is likely to be significantly less than 50%. For instance, BioGRID catalogs at most 1,353,150 interactions between 28,651 unique genetic elements in *Homo sapiens*. This means that, at most (that is, assuming no redundancy between the documented interactions and assuming that interactions are uniformly distributed over genes), 0.16% of possible gene-gene pairs have any documented evidence that BioGRID can scrape from the literature. While BioGRID is an imperfect resource, we believe most experts would agree that a randomly-selected pair of genes have a low probability of demonstrating a quantifiable interaction in a specific cell population such as human dermal fibroblasts.

In our experimental context, however, there is an additional interacting element between the library perturbation and downstream differentially-expressed genes: the lentiviral vector itself. The various stresses that come with pre-transduction protocols, viral transduction, and subsequent drug selection induce widespread transcriptional changes in cells, independent of payload. This represents a significant source of false positives (a gene’s differential expression is mistakenly construed to be due to the library perturbation, rather than experimental stressors), as well as false negatives (either via contradictory effects of transgene and transduction, or due to statistical confounding in the case where the effects of transduction overpower the effects of the transgene).

Consider the case of 55yo cells transduced with GFP, which we treat as a neutral control. We found that 10,536 genes (56.7% of all genes analyzed) were differentially expressed in this population compared to the 55yo control cells. Due to the substantial size of this transduction footprint, we performed two versions of all downstream performance analyses: one with these 10,536 genes included in analysis, and one with these genes excluded. We expect that any “true” performance metrics lie between these two extremes.

To better situate the observed precision of our method – as well as generalize the “mineability” of *quasi* time-series like data beyond tools rooted in nonlinear dynamics – we generate similar gene-gene interaction networks for two very different methods: the statistical method of partial correlations^50,51^ as well as the neural network method DeepSEM.^52^ Additionally, we generate a null distribution of precisions based on a permutation test which randomly generates gene-gene interaction networks. We find that, in all cases, the CCM network has the highest precision (Fig. 2**(e)**). All three methods are statistically highly distinct from the null hypothesis; however, whether DeepSEM and partial correlations are distinctly *better* or *worse* than the null depends on whether the GFP/transduction footprint is included or excluded. In general, this may be due to the advantage of CCM and related methods in inferring causal interactions when the interactions themselves display nonlinear dynamics.^53^

Additionally, we see a broad distribution in precisions among library genes – some library members have nearly perfect overlap between the CCM topology and the differential expression results, whereas some bear no resemblance whatsoever. To best quantify this, we employ a Fisher Exact test, and our metric for *local* CCM network precision is the odds ratio emerging from the contingency table formed from respective overlaps and exclusions of the CCM topology and differential expression results (Fig. 2**(f)**). We find that, in general, the odds ratio per library member significantly improves in the case where the viral transduction footprint was removed from analysis, a trend we continued for the rest of this work. A binary classification analysis on well-mapped versus ill-mapped library members based on available features (see **Methods**) is unable to produce a plausible explanation in terms of the available data – either due to the restricted size of our experimental library, or, perhaps, due to the aformentioned distinction between “essential” interactions that might be discovered by the CCM inference and “aging-specific” interactions that would not necessarily be revealed in our experiments.

As a last measure of network inference performance, we implement a more traditional precision-recall analysis (Fig. S4). This type of analysis varies a key hyperparameter of the network inference: the cutoff CCM strength *ρ_L∗_* (*X, Y*) is used to consider an edge *X → Y* to be valid. The results reflect both on how better to implement the method on this type of data in the future, as well as the extend to which we were over-judicious in our original, pre-experimental choice of cutoff. The results indicate that the optimal network would have been produced using the top 65% of edges, rather than the top 5%.

### Measures of health and rejuvenation in scRNA extracted from perturbed cells

#### Rejuvenative capacity of library members estimated via machine learning

While previous sections focus on gauging the accuracy of the inferred gene-gene interaction network, we quantitatively assess of the rejuvenative capacity of the tested library. In the following sections, we describe and show results for a machine learning approach to quantifying rejuvenation among perturbed cells from scRNA data. We then evaluate the rejuvenative capacity of each library member tested. The results reflect both on this approach to quantifying aging and rejuvenation, as well as the heuristic ranking scheme applied to the inferred gene-gene interaction network for genes with potential ‘rejuvenative’ action in perturbation products.

#### Training a supervised learning ensemble on unperturbed old and young fibroblasts

In order to quantify the rejuvenation induced by gene perturbations on control 55yo cells, we design a supervised binary classifier which takes advantage of our experimental design and the amount of raw data present in single-cell transcriptomics. Because our donor cohort consists of three 55yo and three 25yo donors, with multiple untreated control wells in the scRNA experiment, we treat these as two classes whose transcriptomic ‘fingerprint’ can be established by training a machine learning model to predict the age class of cells selected for each class from untreated wells. The rejuvenative capacity of a library transgene will then be measured by presenting perturbed 55yo cells to the classifier; a high rate of 25yo false positive calls (cells called as 25yo, despite originating from 55yo donors) indicates that a large fraction of perturbed cells have been pushed towards a 25yo phenotype (Fig. 3**(a)**). We employ adaptive boosting (AdaBoost).^54^ AdaBoost is primarily chosen for its sparsity-promoting regularization, resulting in a classifier ensemble that leverages very few genes as features. We benchmark several other methods, including Gaussian-type and SVM-type methods, but never observe competition with AdaBoost in terms of training set accuracy (linear SVM: 51.6%; Gaussian Naive-Bayes: 53.5%).

**Figure 3.**
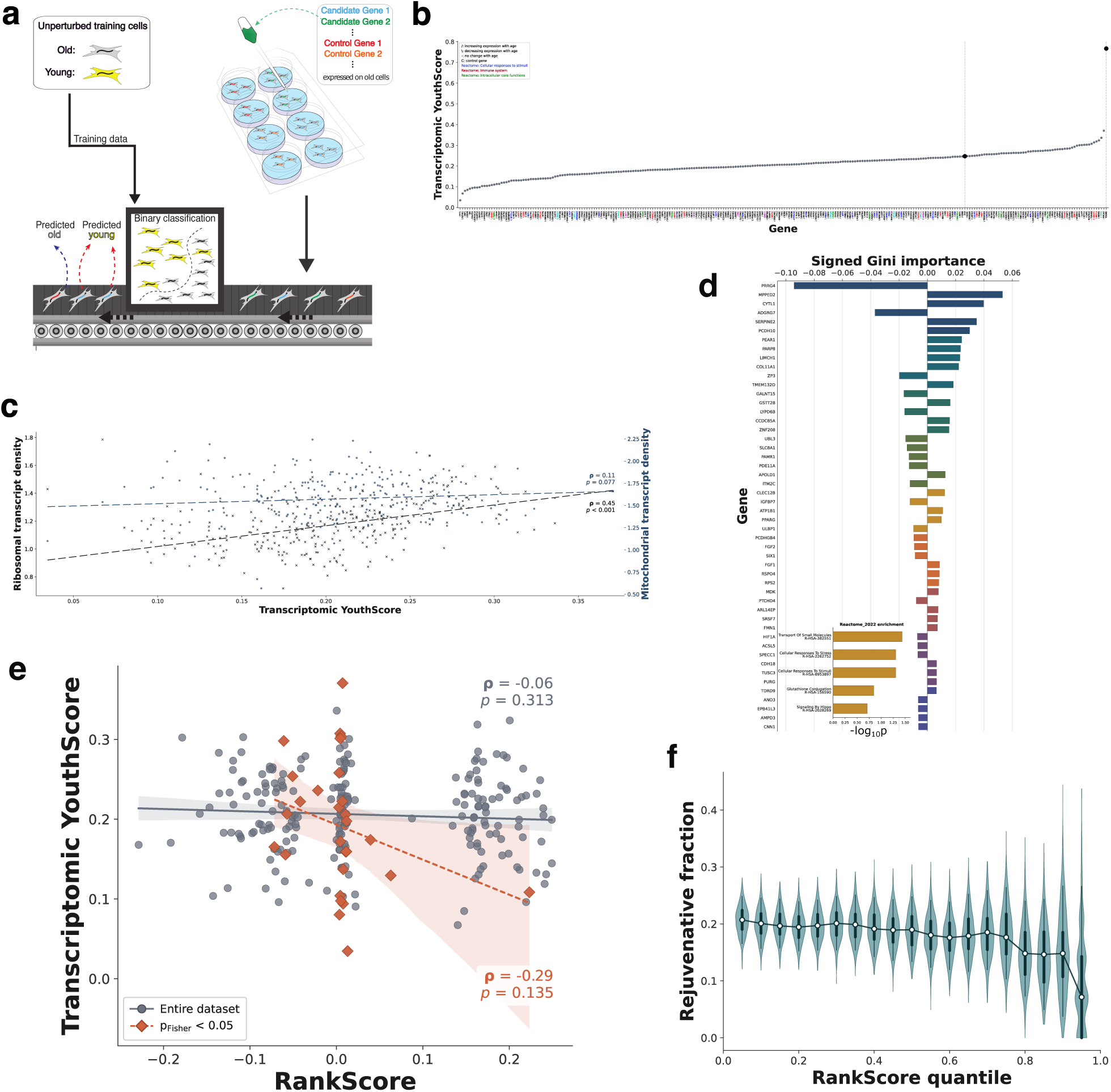
Profiling rejuvenation induced by library member genes, via binary classification. **(a):** A schematic of our approach to quantifying rejuvenation. We employ a binary classification scheme (based on AdaBoost,^54^ see **Methods**), with bagged classifiers for class balance, wherein the training population consists of equal numbers of 55yo and 25yo fibroblasts. The aggregate classifier has a test accuracy of 74.1 *±* 0.7%. We then quantify rejuvenation induced by library gene transduction with the FPR – the fraction of transduced 55yo cells misclassifed as 25yo by the ensemble – which we call the ‘YouthScore’. **(b):** The YouthScore for every library gene, where gene annotations are documented in Figure 2(a). Note that genes which are differentially expressed in GFP-transduced genes have been removed. While most library members are neutral or deleterious, roughly 20% show signs of rejuvenation. **(c):** A quantitative measure of rejuvenation should ideally reflect hallmarks of aging from the literature. Here, quantitative metrics of a healthy metabolic and proteostatic phenotype are rendered in the ribosomal and mitochondrial transcript fractions, quantities typically reserved as QC metrics. Both of these correlate (Pearson) with our YouthScore at statistical significance. **(d):** The genes used by our classifier ensemble, together with their averaged Gini importance. The sign of the Gini importance is positive for genes enriched in 25yo samples, negative for genes enriched in 55yo samples. Inset, we show Reactome enrichment results for these genes. Note that R-HSA-8953897 (“Cellular responses to stimuli”) is a pathway we used to annotate library genes in Fig. 2(a). **(e):** A direct comparison between the ranking score assigned to library genes and the observed YouthScore. For the general library, the association is extremely weak. However, when restricted to the genes with high subnetwork precision (high Fisher Exact odds ratio, Fig. 2(b)), the association increases in strength. **(f):** In analogy to an AURC curve, we plot the fraction of library members seen to have a rejuvenative capacity (YouthScore greater than 55yo controls) when the size of the available library is limited by quantiles of the ranking score. The majority of the rejuvenative gene “discovery rate” (slope) comes from the genes with the highest absolute RankScore. Violins are from 1,000 bootstrapped replicates.

We implement additional measures in our binary classifier to account for the practical challenges of our experimental setup and scRNA data in general. Chief among these is (1) the incorporation of bagging (combining multiple train-test iterations, each with balanced numbers of 25yo/55yo cells) to combat class imbalance due to the differential growth rates associated with aging;^15^ (2) allowing individual bagged models to abstain from voting on the state of test cells. Abstention^55^ is known to increase individual example margins in bagged ensembles, thereby increasing ensemble test accuracy, particularly in cases where the distribution over test examples is heterogeneous. Here, we have both training and test heterogeneity, because our training sets consist of mixed donors, and our test sets (perturbed cells) are highly unlikely to share distributions of features either with one another or with the unperturbed training cells. In these analyses, we omit the viral transduction footprint (differentially expressed genes in GFP-transduced cells, see **Methods**).

Initially, the total ensemble is trained on a standard 70%/30% train-test set drawn from control 25yo and 55yo cells (with class balance enforced). Across bags, the test accuracy is 74.1 *±* 0.7%, with strong consensus on genes of interest (i.e., biomarkers for youth) across models. We find that a relatively low abstention threshold results in an acceptable abstention rate (10% of cells or fewer) with a peak in accuracy boost.

Then, because library perturbations are only applied to 55yo cells, we reason that the response of the classifier to library-member-perturbed cells – both the abstention rate as well as the per-cell-classification – is a quantifiable measure of rejuvenation. Our reasoning is that by rigorously enforcing class balance during training, and additionally allowing abstention, minimizes the possibility that perturbed cells which are “too strange to be 55yo” will be automatically (mis)classified as 25yo. The decision boundary should thus be roughly equidistant from the center of mass of 25yo and 55yo (in RNA space), with a slight “blur” or “no-man’s-land” on both sides of the decision boundary, as well as very far from the training populations. Unnatural cells are very unlike native 25yo or 55yo cells, and so the ensemble will mitigate its log-loss by calling a 50% classification probability for both ages. In addition, to mitigate inherent variability within-donor and across-donor for the perturbed 55yo cells (i.e., to avoid bias in the results due to aberrantly-young 55yo cells falling by random chance into the pool to be transduced by a particular library member), our rejuvenation measure will be determined only by per-cell classification (a binary 55yo versus 25yo), not by class probability.

In this scheme, the measure of rejuvenation is the false positive rate – the fraction of perturbed 55yo cells which are, with some degree of confidence (due to abstention), called 25yo by the classifier. Because the hold-out test accuracy in the training phase is 74%, we know that the “baseline rejuvenation rate” for the control 55yo cells should be roughly 26%, whereas the “full rejuvenation rate” (the effect size which would make the perturbed cells roughly indistinguishable from 25yo cells) is around 74%.

The full range of rejuvenation rates (which we refer to as the “YouthScore”) is pictured in Fig. 3**(b)**. The rates for holdout test controls (unperturbed 55yo and 25yo cells) roughly correspond to the training accuracies (around 25% of 55yo cells misclassified as 25yo, and around 75% of 25yo cells correctly classified as 25yo). We observe that around one-quarter of our tested library members have some degree of rejuvenative effect (a YouthScore greater than untreated 55yo cells). The highest observed rejuvenation rate among perturbations was that of the protein phosphatase PPP1R7, which has a YouthScore of 0.38.

#### Inferred rejuvenation correlates with key phenotypic changes in perturbed cells

Our method for quantifying rejuvenation relies on a key assumption – that machine learning applied to transcriptomic data (genotype) can properly capture changes in cell behavior commonly associated with youthfulness (phenotype). We further test this connection by comparing the YouthScore to phenotypic states.

As our first phenotypic measure, we re-examine the per-cell quantities that are usually taken as quality control metrics: the overall density of mitochondrial and ribosomal reads. Our reasoning is that these measures, especially in perturbation experiments like ours, are subject to a dichotomy: on a per-cell or per-cell-population basis, it is unknowable whether or not changes in mitochondrial and ribosomal count densities (relative to a baseline of control cells) are due to either their intended QC purpose (such as increased mitochondrial count densities indicating damage to the cell membrane), or actual functional changes in the cell (where increased mitochondrial count densities indicate increased mitochondrial activity).

We mitigate the first option (changes induced by damaged or unfit cells), as well as general inter-donor and intra-donor single-cell heterogeneity, by averaging QC metrics across all cells receiving the same perturbation. This scheme is imperfect – a transgene which systematically weakens the cell membrane or depletes cytosolic RNA can still be confused with a transgene that enhances mitochondrial function. However, this seems unlikely to be a general phenomenon in a library of 266 genes. Across library members, we observe statistically significant associations between our transcriptomic YouthScore and both mitochondrial and ribosomal count densities (Fig. 3**(c)**), with the latter association being extremely strong (Pearson *ρ* = 0.45).

Changes in mitochondrial and ribosomal activity are fundamental “hallmarks of aging”,^16^ which are recapitulated specifically in human fibroblasts.^15^ To further cement connections between our YouthScore and other pathways implicated in the aging and rejuvenation literature, we scrutinize the genes which were most important to the entire classifier ensemble (Fig. 3**(d)**). First, we perform a pathway mapping on the *∼* 40 highest-Gini-importance genes based on the Reactome (2022) mapping scheme. We observe enrichment for three common hallmarks of aging: “Transport of Small Molecules”, “Cellular Responses to Stimuli”, and “Cellular Responses to Stress” (Fig. 3**(d)**). Examining individual contributing genes further reveals thematic associations of genes known in the literature. The most prominent of these were well-known shifts in connective tissue and related growth factors, including COL1A1,^56^ FGF1 and FGF2,^57^ and multiple cadherins.^58^ There are also several aging-implicated senescence factors (or youth-related senescence inhibitors), including HIF1A,^59^ APOLD1,^60^ PPARG,^61^ SIX1,^62^ and SRS7.^63^ Finally, we note that synthetic or endogenous increases in at least two of our “marker” genes have recently been strongly implicated in rejuvenation or “blocking” the aging phenotype: EPB41L3^64^ and CNN1.^65^

#### Ranking heuristic on CCM network is partially informative on rejuvenative genes

Our preliminary heuristic ranking (the RankScore, Eq. 1) is chosen for its simplicity, utilizing the most basic level of both dynamic and topological information for each individual gene. As such, we are uncertain to what extent the heuristic would be able to truly predict rejuvenative targets.

The correlation between our RankScore and the experimental transcriptomic YouthScore is pictured in Fig. 3**(e)**. We previously posited that very negative RankScore genes should be the most impactful when overexpressed, since the expression of that gene will generally be decreasing with age. We observe a weak negative correlation between the RankScore and YouthScore. However, we find that the sign of this association changes if the extrinsic (i.e., CCM-topology-driven) component of the RankScore is removed (Fig. S5), which would effectively reverse the logic of whether *over* expression of a gene would represent the correct form of perturbation. Secondly, if we restrict our attention to the library members whose CCM network was found to be particularly precise in our earlier analysis (those genes whose *p*-value from the Fisher Exact test was *<* 0.05) and repeat the analysis, the association strongly increases. However, we note that this newly-thresholded batch represents a small set of genes (40 out of our 266-gene-library).

As a secondary measure of the RankScore’s accuracy, we implement a ranking-based measure. This represents a better measure of the our approach’s practicality, since the absolute value of the RankScore is inherently more subject to noise-driven fluctuations than the ranking based on its value.^66^ Put differently – the applicability of a ranking metric might be better appraised by how often top-ranked items are “successful” than how quantitatively accurate the metric may seem to be.

For this measure, we create a plot analogous to a precision-recall curve (Fig. 3**(f)**). We ablate individual genes in our library based on the absolute value of their individual RankScore. At each value of the ablation, we measure the fraction of tested genes that *would have* come back as rejuvenative (meaning that the observed transcriptomic YouthScore is greater than 55yo controls) in the putative ablated experiment. Beginning with the most-ablated 1-gene experiment (which would have a success rate of zero, as our highest-ranked gene did not induce rejuvenation), the success rate rapidly increases until about 30% of the top-ranked genes are included. At this threshold, the gains from testing more and more genes saturate, until the *∼* 25% success rate observed in the full 266-gene library is recapitulated.

### An orthogonal perspective on rejuvenation via single-cell morphology

#### CellPaint morphological assay on library-transduced cells

While our scRNA results confirm the presence of usable information in *quasi-* time series regarding gene-gene interactions, and our machine learning work on rejuvenation implies that our CCM-based approach may be finding genes with rejuvenative potential, we seek to further validate our approach to quantifying youth and rejuvenation. To this end, we complement our transcriptomic work with a more direct measure of cell health and phenotype using the single-cell morphological assay CellPaint.^67^ CellPaint uses five fluorescent channels to label the structure and activity of five classes of organelles: Hoechst 33342 (DNA), concanavalin A (endoplasmic reticulum), SYTO 14 (nucleoli and cytoplasmic RNA), phalloidin (actin) and WGA (Golgi and plasma membrane), and MitoTracker Deep Red (mitochondria). This assay has previously been used to examine cell phenotypes and cellular responses to drugs,^68,69^ and can extract thousands of features related not only to the basic shape and size of cellular and sub-cellular components, but also correlations and interactions among these components, thereby comprising most features of interest in cell physiology and morphology.

Our CellPaint implementation mirrors our scRNA assay. We use the same mixed three-donor pools of 25yo and 55yo controls, and transduced additional 55yo cells with our *∼* 300 gene CCM-inspired library. We note that, due to differences in experimental protocol, a slightly different fraction of the library is represented in post-quality-control measures than in the transcriptomic experiments – here, 280 library members were represented, compared to the 266 in our scRNA assay (with a 257*/*266 = 96.6% overlap between library members).

#### CellPaint resolves age better than scRNA, and uses more interpretable features

While single-cell RNA sequenciing enables direct evaluation of the CCM results, in addition to the machine-learning-derived quantification of rejuvenation, CellPaint only enables the latter. We therefore directly train our ensemble classifier on untransduced 25yo and 55yo controls. Direct visual examination of the control groups reveals systemic differences, with the 25yo cells having brighter endoplasmic reticulum (ER) images and the 55yo cells generally being brighter in all other channels (Fig. 4 **(a)**). Additionally, the 55yo cells are clearly larger, with larger nuclei and decreased aspect ratio – all known phenomena in fibroblast aging.^70^

**Figure 4.**
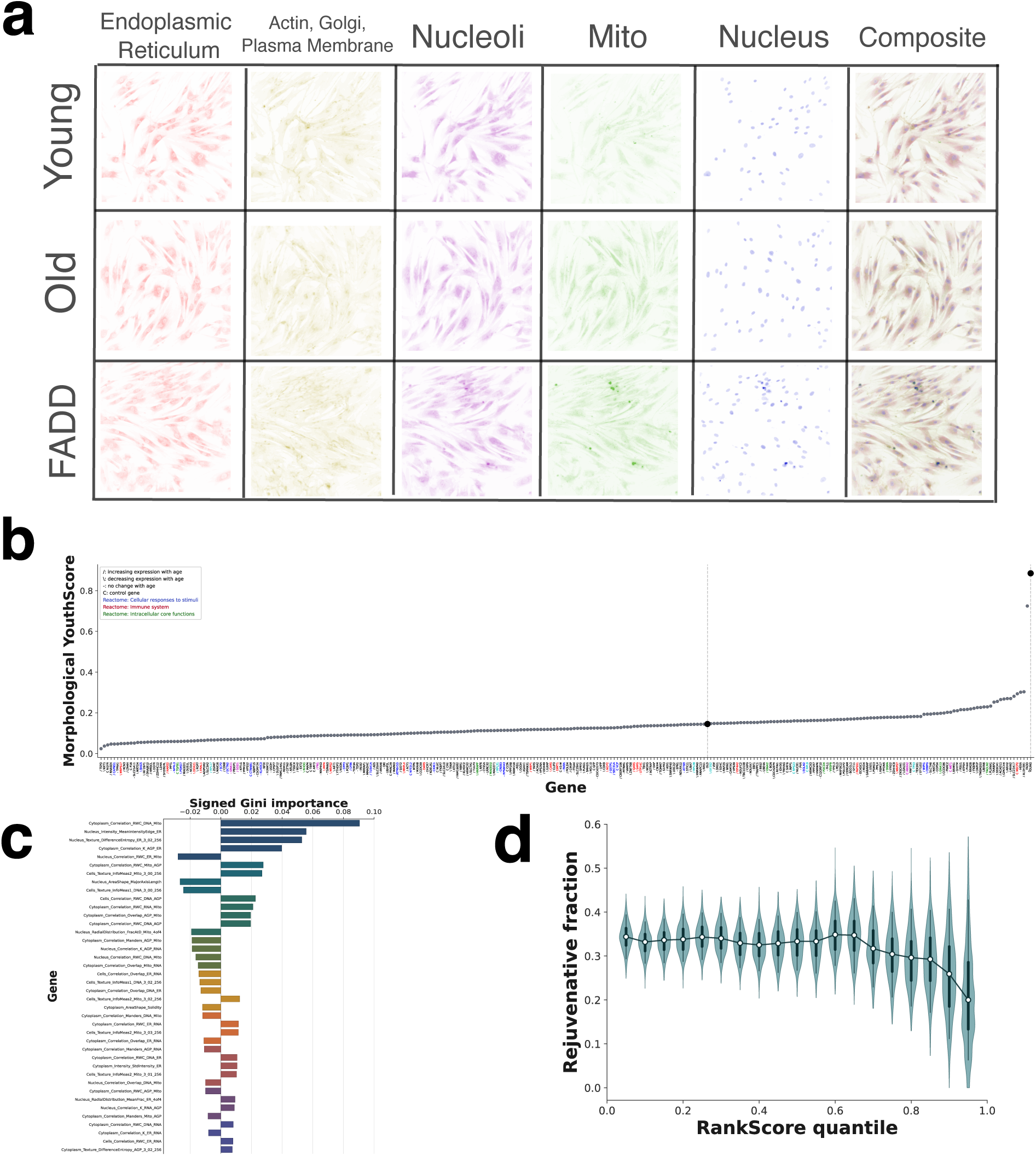
Morphological profiling via CellPaint assay. **(a):** CellPaint is an imaging technique utilizing five stained channels to identify markers related to the endoplasmic reticulum (ER), actin/Golgi/plasma membrane (AGP), DNA, RNA, and mitochondria. This example shows all five channels (and their combination) for dermal fibroblasts taken from pooled 25yo donors (top row) and pooled 55yo donors (bottom row). **(b):** A morphologically-derived YouthScore, which shows a wider spread than its transcriptomic counterpart (Fig. 3(b)) in terms of effect sizes, with evidence of stronger rejuvenative effects (with the best library member, FADD, being “extremely close” to the true 25yo control cell population. **(c):** Compared to scRNA, the features driving the high accuracy (83.1 *±* 0.5%) of our ensemble classifier have higher human interpretability and are easier to correlate with known biology from the aging literature. **(d):** The analogy of Fig. 3(f), again recapitulating that the highest discovery rate of rejuvenative genes comes from targets with the highest absolute RankScore.

The ensemble classifier picks up on this basic physiology, as well as differential growth rates between old and young cells. The most informative features of cell age (Fig. 4 **(c)**) include larger sizes for older cells (negative coefficients for *Nucleus AreaShape MajorAxisLength* and *Cytoplasm AreaShape Solidity*). The differential growth rate, as represented by the the distance between nearest neighbor cells, was a feature with very high Gini importance, leading us to remove it from the feature set in order to identify subleading features associated with the age state (see **Methods** for more details on feature engineering).

The CellPaint-driven classifier ensemble showed markedly higher accuracy than the scRNAseq results: 83.1 *±* 0.5%, roughly 10% higher. This matches intuition that CellPaint features more directly match the cell’s“true phenotype” than transcriptomic features.

#### CellPaint rejuvenation measure shows larger effect sizes than scRNA

Given the increase in classifier accuracy with CellPaint features we also compute a morphological YouthScore (Fig. 4**(b)**), and we observe a larger spread in rejuvenative effect sizes compared to our transcriptomic YouthScore. The overall fraction of rejuvenative targets is higher (around 36% for CellPaint versus *∼* 25% for scRNA), and a significant fraction of rejuvenative targets have YouthScores which are at least twice that of 55yo control cells, a feat not achieved by a single library member according to the transcriptomic measure.

One particular target, FADD, has a morphological YouthScore which was nearly four times the 55yo control baseline, with its cells called (erroneously) by the ensemble as 25yo only 10% less frequently than genuine 25yo cells. This particular target is one of the small fraction that failed quality control in our scRNA experiment, with only a handful of cells surviving drug selection.

In comparison with our CCM results, we observe a correlation near statistical significance (Pearson *ρ* = *−*0.11, with a *p*-value of 0.06). However, this correlation still shows a weak overall effect size. We performed a similar “hit rate” analysis, finding again that the fastest discovery rate of rejuvenative genes comes from the genes with the highest absolute RankScore (Fig. 4**(d)**).

#### CellPaint and scRNA YouthScores are strongly correlated, implying a coherent approach to quantifying rejuvenation

To cap off our work, we restrict all available data to a set of genes which appear in both the transcriptomic and morphological experiments. Then, we examine the Pearson correlation between a number of relevant features, including our scRNA QC/phenotypic measures, both our transcriptomic and morphological YouthScores, our CCM-driven RankScore, and our *post hoc* analysis of the CCM network accuracy (Fig. 5**(a)**). Here, we restrict our attention only to those that show statistical significance (*p* ⩽ 0.05 after multiple-hypothesis correction).

**Figure 5.**
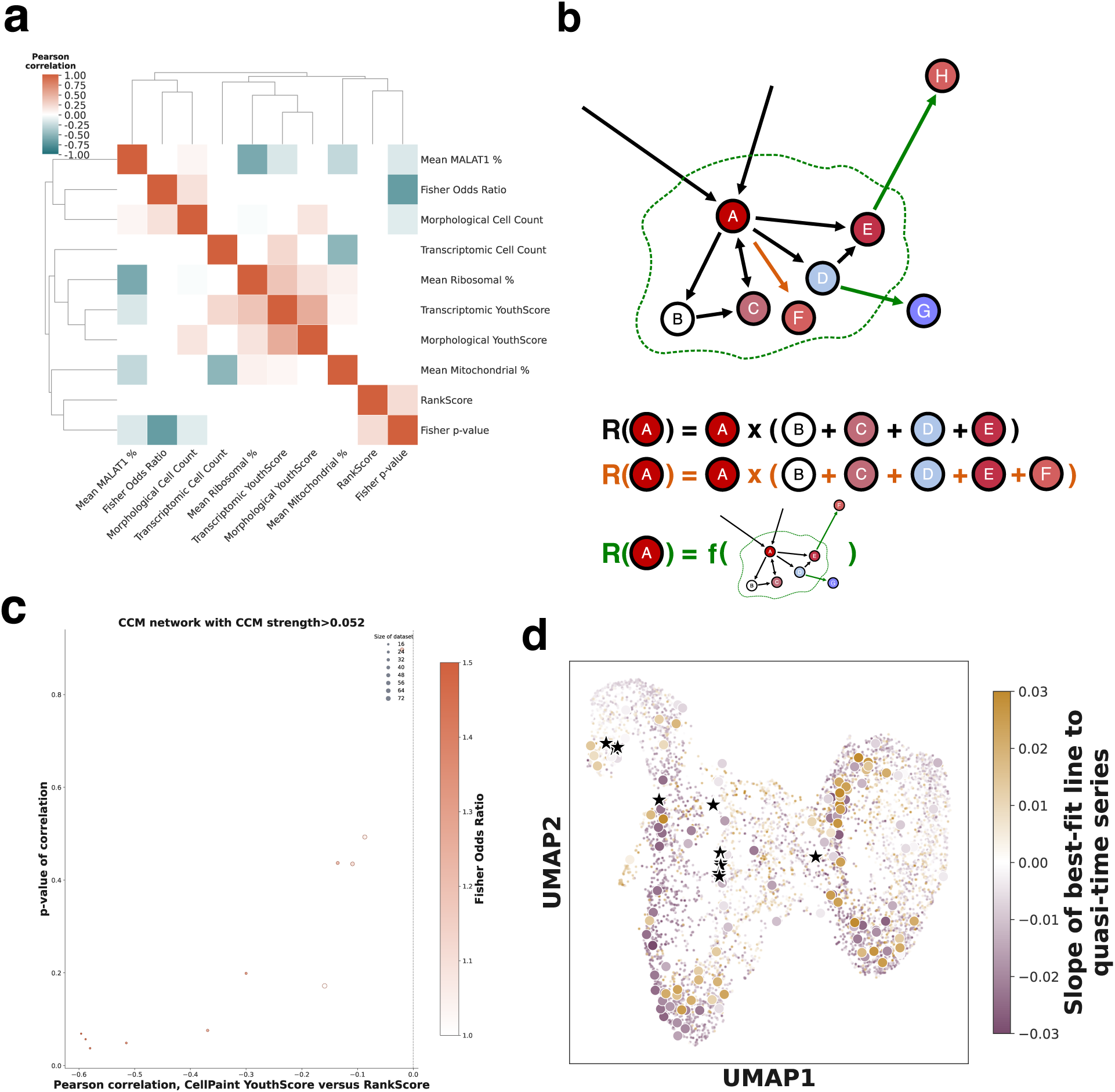
A comparative survey of results, and a loosening of base assumptions. **(a):** A (Pearson) cross-correlation matrix between features derived from all areas of our analysis, restricted to a set of cells / library members which were present in both the transcriptomic and morphological analyses. The strongest correlation (Pearson *ρ* = 0.52) is between the independently-derived transcriptomic and morphological YouthScores. Other interesting correlations include the YouthScore-Cell Count correlations for both transcriptomic and morphological studies, as well as the RankScore positively (not negatively) correlating with the *p*-value from the Fisher exact test on CCM “well-mappedness”. **(b):** Search space for improvements to the RankScore. We consider two principal axes of extended search for a RankScore that more closely correlates with the experimental YouthScores; the first involves augmenting the CCM topology used throughout the manuscript and implementing the base RankScore heuristic (orange), whereas the second axis goes beyond the one-hop nature of our base RankScore heuristic to include deeper probing of network topology. **(c):** Through using larger network topologies arising from looser cutoffs *ρ\** in the CCM network, we observe improved results compared to those found in the main text. Here, the network that is found to be optimal in terms of precision (see Fig. S4) gives an extremely strong and significant match between RankScore and (CellPaint) YouthScore. **(d):** We train a Node2Vec model of our CCM network (see **Methods**), which embeds the network topology – including multi-hop paths – into a low-dimensional space. Using the Node2Vec coordinates, CCM-strength-weighted edges, and the node weighting employed in the heuristic base RankScore, we achieve a better-fitting model than our base RankScore which has additional predictive power. Here, we show a UMAP projection of this feature space; points represent genes in the CCM network, colored by the node-weighting scheme from the base heuristic. We highlight tested library members with rejuvenative effects by vastly increasing their size. Finally, black stars indicate twelve genes which we did not test, which are predicted by the Node2Vec model to be rejuvenative.

The significant results of this cross-correlation analysis reveal a number of interesting statistical/experimental as well as biological relationships. Some confirm known effects, justifying our technical and methodological choices. For instance, the usage of MALAT1 count densities as a QC feature is supported by the fact that it anticorrelates with mitochondrial and ribosomal count densities as well as transcriptomic YouthScore.

Other revealing relationships appear, beyond these sanity checks. For instance, both the transcriptomic and morphological YouthScores correlate with the overall abundance of cells observed to be transduced with library members in each experiment. This could admit multiple interpretations – in one sense, it could simply be recapitulating the known fact that younger fibroblasts divide faster, supporting our YouthScores as accurate measures of rejuvenative library member effect. It also may indicate the role of sample size in establishing the YouthScore. If a sufficiently small population of cells are transduced with library member, then the effects of transgene – whether beneficial or deleterious – may not fully manifest themselves in the population-wide YouthScore measure.

We believe the most impactful cross-correlation, however, to be the strongest one observed – that between the morphological and transcriptomic YouthScores (Pearson *ρ* = 0.52). Given the differences in overall effect sizes, classifier ensemble accuracy, and the incidental fact that some high performing genes (such as FADD) are not present in the post-QC data matrices for both experiments, make this relationship non-obvious until directly measured. The correlation between two completely different length scales of information – the nanoscale of RNA, and the microscale of cell physiology – supports the idea of holistic genotype/phenotype “fingerprinting” of age and youth, as well as rejuvenative interventions.

We next performed a literature search on genes which we found to have a rejuvenative effect in both our scRNA and CellPaint work (Table S1). In particular, we qualify to what extent these genes have been investigated in the context of aging, age-related disease, and/or rejuvenation; we were also interested in qualifying the potential pathways or mechanisms-of-action involved in our observed effects. Overall, the distribution of rejuvenative genes mimics the distribution of our full tested library in terms of RankScore behavior as well as the frequency of transcription factors, surface markers, and secreted factors (15% in rejuvenative genes, compared to 13.5% in the total library). Roughly half of our rejuvenative genes are implicated in the aging literature, though this is a subjective matter; in the citations provided, we restrict ourselves to studies focused on mammalian tissue (though we note genes that are implicated, for instance, in *C. elegans* and *Drosophila*, where lifespan studies are much more robust), and studies where the focal gene is a prominent subject of the study (for instance, appears in the manuscript’s primary claims, rather than belonging to large lists of differentially-expressed genes). In terms of pathway/mechanism, we find general bias towards mitochondrial/metabolic activity, and the *p*53 constellation of pathways.

#### Attempting to find *post hoc* improvements on the RankScore

Our experimental results allow us to investigate alternative formulations of our RankScore, in addition to validating it. All the choices involved in designing our CCM and downstream inferences affect the downstream RankScore, even when implemented in its heuristic formulation (Equation 1); furthermore, it is clear that the shorthand heuristic nature of this base RankScore could be expanded upon greatly. Here, we consider two directions for refinement: firstly, through keeping Equation 1 as the foundation of the RankScore, but expanding or tweaking the topology of the underlying gene-gene interaction network; secondly, through considering extensions of 1 in quantifying the RankScore. These two approaches are schematized in Fig. 5**(b)**.

We have observed a few indicators in this work that the primary CCM topology used and the base formulation of the RankScore do not interact optimally. This weakness, which would have been difficult to detect before performing experiments, primarily hinges around how the RankScore behaves for well-mapped and ill-mapped gene neighborhoods according to our scRNA screens and associated Fisher Exact results for CCM accuracy. The most succinct proof of this weakness is the positive correlation in Fig. 5**(a)** between the RankScore and the Fisher Exact *p*-value. The direct meaning of this is clear – the higher the RankScore, the higher the Fisher Exact *p*-value, meaning that the neighborhood of the focal gene in both cases was ill-mapped. However, the interpretation of this correlation is interesting due to the signedness of the RankScore. The general trend here means that very negative RankScores – which we have loosely interpreted as corresponding to prime targets for rejuvenation-via-overexpression – may also have been subject to more precise CCM inference. The temporal dynamics of each gene in the network should therefore also be included in any consideration of possible improvements to the RankScore.

The simplest change to implement and examine is to consider alternative network topologies. Here, we retain our CCM-inferred network, and change the topology solely through manipulating *ρ\**, the cutoff end-library CCM strength above which two genes will be connected with an edge. There are two natural alternatives to our initial, stringent choice of *ρ* >* 0.5677; these are *ρ* >* 0, which would admit as a gene-gene interaction any pair of *quasi* - time series which have some predictive power at full data availability and which show an increase in this predictive power as a function of data availability. The second choice is *ρ* >* 0.052, a threshold which we discovered in our scRNA-fueled *post hoc* analysis of the CCM network, which represents the most precise topology available to us given present information (see Fig. S4). For both of these topologies, we re-calculated the RankScore according to Equation 1, and re-performed correlations between these RankScores and the measures of rejuvenation derived from experiment.

We saw massive improvement in the relationship between RankScore and YouthScore (particularly the CellPaint-derived YouthScore) in the *ρ* >* 0.052 topology (Fig. 5**(c)**). This statistical relationship kept its sign, but the effect size increased and statistical significance (*p*-value less than 0.05) was attained in several cases, especially when the genes under consideration were restricted to those whose CCM network is considered to be well-mapped. See Fig. S8 for a similar treatment of the *ρ* >* 0 topology, as well as the scRNA-derived YouthScore.

These improvements are heartening, as they suggest that direct improvements to the accuracy of the CCM topology would lead to more efficient discovery of rejuvenative genes – arguably the core thesis of our work. However, they still rely on the base RankScore heuristic, which was intended only as an initial “guess” at how dynamics and CCM-inferred topology might interact at the timescale of aging to drive key phenotypic changes. Natural extensions to this RankScore are extensions of the underlying feature space, both the intrinsic dynamic component and the extrinsic interaction-topology component.

We explore both simultaneously by considering different intrinsic and extrinsic featurizations, and abandoning Equation 1 as the main quantitative ranking of these featurizations. As a replacement for Equation 1, we explore a broad class of statistically-derived models (since we can regress their predictions onto the experimentally-derived YouthScore(s) to refine model selection). Extensions to the feature set for each gene include alternative dynamical quantities which can be extracted from the *quasi-* time series, as well as deeper probing of the network topology (that is, longer walks, rather than the one-step heuristic of the base RankScore). Deeper sampling of the network can be worked into the feature set for each gene by considering its position in a low-dimensional graph embedding of the larger topology, here implemented via Node2Vec (see **Methods**).

Casting a wide net in terms of combinations of statistical model classes and featurizations leads us to a pleasing optimum – a relatively simple and sparse LASSO^71^ model, based on the smallest training gene set and featurizations explored. This gene set is the well-mapped genes by the Fisher Exact test, featurized solely by the dynamics-based node weighting (the ‘heat’ pictured in Fig. 1**(g)**) and the Node2Vec topological features. This system is comparable in structural simplicity to our base RankScore, but with the one-hop step replaced by a deeper probing of the topology; it also shows a stronger association between the imputed RankScore and the experimental YouthScores than those we describe in the main text, and the statistical significance is underscored by strong performance across multiple holdout cross-validation test sets.

These refinements to the RankScore show how the programme we lay out in this work could be viewed as a self-improving, self-refining loop. Consider Fig. 5**(d)**, which shows a UMAP representation of the feature set used in the Node2Vec-driven model. We have highlighted the genes explored in this study which had a rejuvenative effect, which show an unusual clustering pattern across the UMAP embedding. Additionally, we highlight the ten genes with the highest Node2Vec-derived RankScore which were *not* included in our experimental library – while most of these cluster near the tested genes, some are relatively isolated. Positive results emanating from future examination of these genes could lead to yet another refinement of the gene-gene interaction network inference and rejuvenation target ranking, the first step in an iterative program.

## Discussion

Aging is perhaps best described as a complex of changes that take place in an organism at all scales as a function of “long time”, to be distinguished from shorter timescales such as circadian and seasonal oscillations. While this is a vague description, it has the advantage of being “hypothesis-free”; it does not assume that any hallmarks of aging have causative effect, nor does it assume that drivers themselves will emerge as leading biomarkers or predictors of aging. The idea of “long time” also obscures the underlying biology, because the nature of experiment biases us towards a greater understanding of the dynamics that take place on shorter timescales. This means that the extent to which well-known interaction mechanisms such as transcription factors and RNA interference also affect aging is essentially unknown. We believe this necessitates a more agnostic, data-driven approach to understanding how various -omics quantities might interact in concert to generate the aged phenotype(s).

Our *quasi-* time series composition and computational inference approach has the advantages of being agnostic, inductive, and adapted to the dynamic nature of aging data. With comparatively meager data – equivalent to seventy datapoints per gene, readily available in public databases – this approach discovers gene-gene interactions which are broadly supported by literature evidence and by experiments probing short-timescale transcriptomic dynamics. We observe that this approach is also extremely computationally efficient, and could therefore easily be scaled to many more gene species and tissue types. While this is not intended as a benchmarking study, our choice of inference method outperforms selected comparanda methods by these measures. Even though our goal is a broader understanding of aging, the fact that our validation methods all rely on short-time biology suggest that this approach could be useful in discovering short-time gene-gene interactions which are preserved in the long-time dynamics of aging. Conversely, it is also possible that the metrics of our method’s accuracy have been lowered due to the existence of long-time biological dynamics that do not manifest themselves strongly enough (or at all) on short-timescales to be adequately detected.

We believe that these inferred gene-gene interaction networks have predictive power beyond transcriptomic interactions, and can be used to suggest genes of interest that might have the ability to alter or reverse the aged phenotype when their expression is perturbed. To this end, we develop our transcriptomic and morphological YouthScores, machine learning models that show high training accuracy on controlled aging cell models and which theoretically should have strong generalized error bounds. These models yield binary age classifications for single cells that correlate strongly with known hallmarks of aging, such as basic densities of mitochondrial and ribosomal RNA molecules. We use this as our preferred marker of the putative aging and rejuvenative activity of a gene, reasoning that a cell that “looks young” on a holistic transcriptomic and/or morphological scale can be equated, roughly, with a genuinely chronologically young cell. Our morphologically-derived YouthScore showed a greater training accuracy and downstream effect sizes than our transcriptomically-derived YouthScore, perhaps in line with a Central Dogma-derived argument that cell physiology and morphology should be better predictors of cell age and cell health than transcriptomic activity. Nonetheless, the two YouthScores generally correlate with one another in addition to known hallmarks of aging. In both the transcriptomic and morphological experiments, genes of interest identified from our interaction networks produced significant rejuvenation according to our YouthScores. In one particular instance (the gene FADD, which appears in our morphological experiments but failed to produce enough viable cells post-drug-selection in our transcriptomic experiments), the degree of rejuvenation seems to make the majority of 55-year-old cells nearly indistinguishable from 25-year-old cells.

These results suggest that there is value in applying modern computational methods – especially those that incorporate aspects of dynamical systems theory – to extracting biological knowledge from aging data. Furthermore, that aging data need not be proprietary data which is expensive to generate or obtain; even publicly-available, bulk RNA measurements, concomitant with all the noise that comes from large population samples, contain valid and actionable signal. The mathematical foundation of our program suggests that this actionable signal derives from the fact that gene expression in long, aging timescales shows signs of *recurrence*. Rather than linear trends of expression with age, which dominate the aging biomarker literature and which are intuitively easier to grasp, we find that the vast majority of our aging *quasi* - time series show cyclical and recurrent behavior. This is supported by other studies in recent years, which show statistically recurrent and even periodic time courses for various -omics quantities at the aging timescale.^72–74^

We intend our study to be the first validation step in a broader program to extend our understanding of aging dynamics, and as a consequence there are a number of shortcomings and outstanding questions. We find that our CCM-based inference approach shows a wide spread of accuracies at different points in the inferred interaction network topology, at least when accuracy is measured by short-time biological experiments. We are not able to account for dynamical or biological causes for these differences in accuracy. They could arise from different distributions of noise in the underlying *quasi* - time series, or in sources of inductive bias in our methodology that we are not able to pinpoint. The spread of CCM accuracies has a significant impact on our downstream analyses; we observe very different correlations between prediction and experiment based on whether the whole CCM network or various CCM subnetworks based on observed accuracy were employed. Even in the best cases, however, our attempts to predict rejuvenative genes from our *quasi-* time series were imperfect. Our results suggest that the program is a significant improvement on a random experimental library, but we believe that there are a number of avenues worth pursuing to refine this approach. The most obvious barrier to improvement within our own study is the limited experimental library. Repeating these experiments with a broader library would significantly improve the ability to determine what dynamical trends make a gene worthy of focused experimentation.

There are also a number of technical choices in our study that we believe deserve further scrutiny and represent interesting research directions on their own. Our method of generating *quasi-* time series, which involved averaging over all donors of the same age, likely represents the absolute simplest possible method of collapsing population-wide single-measurement bulk RNA data into a time series format. Considering alternative methods of generating these time series may not only furnish higher downstream performance, but also genuine biological insight even absent downstream analyses. For instance, methods of extracting *quasi-* time series that incorporate the theory of “ageotypes”^11^ could quantify the abundance and diversity of these trends in population samples. Our machine learning AgeScores are also novel, and we deliberately used them in place of other methods which exist in the literature, such as “aging clocks”. We made this decision based on technical considerations, and the design of our experimental studies does not offer the primary data which would be necessary to rigorously establish connections between our YouthScores and other computational efforts to quantify aging and rejuvenation. We believe investigating these questions could offer not only biological insight, but a deeper understanding of how machine learning and data science methodologies are best implemented in aging biology.

## Methods

### Generation of *quasi-* time series

The empirical data used to infer gene-gene interaction networks was taken from the “Cells – Cultured fibroblasts” subset of GTEx (V10), consisting of dermal fibroblasts from 490 donors with ages ranging from 20 to 70. 173 (35.3% of donors were female); 2 self-identified as American Indian, 5 as Asian, 60 as Black, and 3 as Unknown, leaving 420 (85.7%) as White. The target sequencing depth (bulk RNA) for each donor sample was 50 million reads.19 The original alignment library (GRChv38) contains 56,200 distinct genetic elements. However, most of these sequences read at zero or near-zero counts in most or all donors; we therefore began by retaining only the 5,991 genes which had nonzero reads in more than half of the donors and which additionally had *zFPKM > −*3.^75^

The raw counts provided in the GTEx data were then normalized as transcripts-per-million (TPM), where the transcript lengths were taken from annotation files belonging to the same alignment library (GRChv38). Because our principal analysis (CCM, described below) is a physics-based (alternatively, “dynamics-based”) approach, we believe TPM normalization to be preferable to counts-per-million (CPM) normalization. This is because the additional step in TPM normalization which scales observed counts by the length of the generating transcript is attempting to capture a physical observable from within a cell/sample, whereas CPM is more of a statistically-oriented normalization.

Finally, we generated the *quasi-* time series for each gene abundance as follows: each donor *j* is represented by a vector *v_j,g_* of expression levels of all genes *g* in TPM. The donor additionally has an associated age *i*, measured in years. A *quasi-* time series for all genes in the dataset will be a matrix *M_ig_* with one expression level per gene *g* and per year *i ∈ {i_min_, i_min_* + 1*, …i_max_}*. If there is ⩾ 1 donors in the set *j\** matching the age index *i*, then the *quasi-* time series entry will simply be the average over those donors: *M_ig_* = Σ*_j∗_ v_j,g_/||j*||*. If *j\** is empty, then *M_ig_*is generated via linear interpolation.

### Convergent cross-mapping (CCM)

For each of the measured 5, 991 genes, we computed the interaction strength with all other genes via Sugihara *et al.’s*^14^ convergent cross-mapping (CCM) approach. CCM accomplishes this by transforming an important theorem in dynamical systems theory into a leave-one-out cross-validation test for mechanistic coupling. Takens’ theorem^28^ states that if two variables *X* and *Y* are coupled in a dynamical system, they have the property that, when embedded in the variables (*X*(*t*)*, X*(*t −* 1)*, …X*(*t − E*)) (a *delay embedding of order E*; same embedding for *Y*, so that both variables are embedded completely independently of one another), local neighborhoods are preserved – meaning if the nearest neighbors in the *X* embedding of observation *X*(21) are *{X*(3)*, X*(7)*, X*(192)*}*, then if *X* and *Y* are coupled, the nearest neighbors of *Y* (21) should be *{Y* (3)*, Y* (7)*, Y* (192)*}*. This can be rigorously tested via the following outline:

1. Create the order-*E* delay embedding of *X*(*t*) and *Y* (*t*) (in this work, we tested *E ∈* [3, 4, 5, 6, 7, 8, 9, 10]), using a portion *L\** of the total length of the time series (*L*).
2. To test the idea of a mechanistic coupling *X → Y*, discard a single observation *X*(*t\**) (because if *X → Y*, it is *Y* that contains information about *X* and not the converse); the discarded datum will be approximated using data from *Y*).
3. The *E* + 1 nearest neighbors of *Y* (*t\**), *{Y* (*t*_1_)*, Y* (*t*_2_)*, …Y* (*t_E_*_+1_)*}* are identified (because *E* + 1 observations are necessary to triangulate a point in *E*-dimensional space).
4. *{X*(*t*_1_)*, X*(*t*_2_)*, …X*(*t_E_*_+1_)*}* are interpolated to produce an approximation *X*^^^(*t\**) for the discarded observation.
5. Repeating this for every *t\** produces a complete time series *X*^^^(*t*), which is an approximation based on assuming *X → Y* holds and using Takens’ delay embedding. The Pearson correlation between the real time series *X*(*t*) and the approximant *X*^^^(*t*), *ρ_L∗_* (*X, X*^^^), is measured.
6. The process is repeated for all *L\** ⩽ *L*. This gives a relationship between how good the Takens’ approximant is, (*ρ_L∗_*), versus how much data was used, (*L\**). The relevant measures from the test are *ρ_L_*, the strength of the test with the most data included, and also the Spearman correlation *corr_Sp_*(*L*, ρ_L∗_*) between *L\** and *ρ_L∗_*, because in a true coupling the slope between the amount of data we have and the amount of predictive power we have should be monotonic.

For each of the 5, 991 *×* 5, 990 = 35, 886, 090 (self-maps are redundant) total possible cross-maps, we calculated the cross-mapping skill as a function of library length *L* (which cannot exceed 51, the total number of measurements) and embedding dimension *E*. We then performed Spearman rank-correlation on cross-mapping skill and *L* for each cross-mapping to isolate those that showed convergent cross-mapping (by keeping only those with *p_Spearman_*⩽ 0.05). This narrowed the search space to 11.5 million edges. Because the distribution of CCM strengths at maximum library size (*ρ\**) is normally distributed (see Figure S1), we decided to keep the top 5% of interactions ranked by *ρ\**, in analogy with the statistical *p*-value. This resulted in a minimum *ρ\** of 0.5677, and a network of nearly 1.8 million interactions between 5,162 genes. In what follows, this directed network is often represented by its (non-symmetric) binary adjacency matrix Δ*_ij_*, consisting of values of 1 if a directed edge from gene *i* to gene *j* was inferred, and 0 otherwise.

### BioGRID statistics

Our quantification of the frequency of BioGRID literature matches for our CCM-inferred interactions came directly from the BioGRID API. The BioGRID build used was 4.4.238. This was implemented in Python, using HUGO gene names, following the code provided in the API documentation. Other stated figures, such as the size of the database (approximately 1.34 million documented interactions, with only 0.1% of the possible 28, 594 *×* 28, 593 gene-gene pairs having any documented interaction) are taken from the build statistics.

### Meta-analysis of the CCM results

At certain points in the manuscript, we attempted to probe the inductive bias of the CCM network by examining how featurizations of a gene (based on both the dynamics of the *quasi-* time series and the inferred network topology) could predict our results, namely, why some the neighborhood of some genes in the CCM network had very high or low precision in our differential expression analysis, and how higher-order ranking heuristics could squeeze more information out of the raw data and CCM network.

To featurize the dynamics of each *quasi-* time series, we used the **extract features** command from the tsfresh Python package with default hyperparameters and settings on the TPM-normalized *quasi-* time series. This returns 783 features per gene / *quasi-* time series. The features cover several concepts from dynamical systems theory, such as the moments of the distribution of the *quasi-* time series as well as its first difference; information-theoretic measures such as the entropy of the *quasi-* time series, its first difference, and relevant transforms such as the Fourier transform; signal-processing measures such as the energy in relevant transforms; and other metrics.

The featurization of the CCM network itself – including nodewise properties such as in-degree, out-degree, as well as nodewise and global properties used later in the work such as nodewise centralities – was performed using the Python package networkx on the adjacency matrix generated via the described CCM procedure.

### RankScore calculation

While there are many possible metrics within the scope of our data which could predict the relative importance of a gene in the aging process, we wanted something which took advantage of the network structure emerging from the CCM analysis and which would have some degree of robustness against the inevitable amount of noise in the underlying dataset.

Additionally, we reasoned that a measure of a gene’s importance in the aging process should take into account: (i) the amount of change-in-expression observed in the gene itself, and (ii) the amount of change-in-expression observed in genes which are downstream from the gene in question in our inferred directed network given by the adjacency matrix, Δ*_ij_*. To factor in some amount of resistance to noise, we quantified “change-in-expression” in a manner that incorporated the whole *quasi-* time series globally. We took the *p*-value of the Spearman rank correlation between the data *M_ig_* and the time axis (*i*), representing the degree to which *M_ig_*is monotone (as well as implicitly quantifying the amount of noise/uncertainty in “change-in-expression”, since a volatile *quasi-* time series will inherently have a higher *p*-value). This measure of monotonicity is then multiplied by the slope *S* of the best-fit line to the *quasi-* time series. The “change-in-expression” for a gene *g* is then *S_g_*(1 *− p_g_*) – if a gene changes very little with age, *S_i_*will be small; if it changes quite a bit, but is also quite volatile and therefore possibly unreliable, (1 *− p_i_*) will be small. The importance, or “RankScore”, is then the amount of change of one gene times the sum change in all the genes receiving a directed edge from the focal gene:

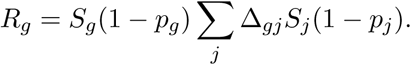

We also considered an alternative scheme, which we comment on at certain points in the paper. In the above expression, *S_j_* – the year-to-year rate-of-change in the expression of a gene *j* downstream from the focal gene *g* – could be positive or negative, with the number of genes with a negative slope generally outnumbering those with a positive slope (though the TPM normalization puts constraints on the global slope behaviors, since Σ*_g_ M_ig_* = 1 *∀i*). This primary RankScore will therefore be small for focal genes of positive and negative change-in-expression. The alternative scheme removes this possibility by instead considering only the absolute value of the change-in-expression which is “induced” by the focal gene:

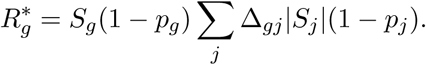

### Reactome querying

In many of our figures, our experimental library of *∼*300 genes (266 in our scRNA screen versus 280 in our CellPaint screen) is annotated with a number of key pathway mappings. The annotation scheme, ultimately, is that of Reactome. However, the analysis used to determine the association of a library gene with one or more pathways was performed using the GSEAPY package and its Enrichr API. The Enrichr version of Reactome used is “Reactome 2022”. A gene was associated with a pathway if the associated *p*-value (from a Fisher Exact test comparing gene inclusion/exclusion in the pathway, with no background set of genes specified) was *p <* 0.05. No multiple-hypotheses correction scheme was used for these *p*-values, as they are ultimately representing qualitative associations. In many cases, the precise name of the mapped pathway is quite technical (i.e., narrow in scope) and differs from our listed pathway name. However, since the Reactome pathways (like many annotation schemes) have a hierarchical structure, we freely combined associations with sub-pathways into three larger super-pathways in order to facilitate easy visualization. While the inclusion (and, comparably, exclusion) of certain super-pathways in both our experimental library and the broader CCM network is intriguing, we did not observe a sufficiently strong correlation between any super-pathway and rejuvenative activity to merit “rigorizing” the described pathway mapping procedure further.

### scRNA experimental details

#### Cells

Human dermal fibroblast cultures were purchased from the Coriell Institute for Medical Research. For the 25yo cell populations, 3 male donors and 3 female donors with an average age of 25 years were pooled at equal cell counts. The 55yo donor population was constructed in the same way, but with donors having an average age of 55 years. All cells were cultured in Dulbecco’s Modified Eagle Medium/Nutrient Mixture F-12 (DMEM/F12) supplemented with 15% Fetal Bovine Serum and 1% Penicillin/Streptomycin at 37°C with 5% CO2 and 95% relative humidity. Cells were maintained at a density between 50-90%, and all experiments were performed with cells at *<* 7 passages.

HEK293T cells were purchased from ATCC. The cells were grown in Dulbecco’s Modified Eagle Medium (DMEM) supplemented with 10% Fetal Bovine Serum and 1% Penicillin/Streptomycin at 37°C with 5% CO2 and 95% relative humidity. Cells were kept under a density of 85% and all experiments were performed with cells no later than 17 passages.

#### Lentiviral vector construction

The lentiviral entry collection was purchased as a clonal plasmid library in plasmid and glycerol forms from Transnomics. The collection was purchased as individual clones arrayed in 96 well plates.

To create our screening collection, genes of interest were cherrypicked using Bravo liquid handling robot in the ICCB-L at Harvard Medical School. Each clone was sequence verified by Sanger sequencing. For controls, the following were custom cloned and LR’ed into the pLX-304 destination vector (identical to the backbone of our screening collection).

#### Lentivirus production

A total of 2.56 *×* 10^4^ HEK293T cells were seeded into each well of a 96-well plate in Dulbecco’s Modified Eagle Medium (DMEM) supplemented with 10% fetal bovine serum (FBS) and 1% non-essential amino acids (NEAA). After 24 hours, each well was transfected with a mixture containing 0.192*µL* of PEI transfection reagent, 10*µL* of Opti-MEM Reduced Serum Medium, 28.2 ng of the Orfeome gene of interest, and 35.8 ng of the Cellecta Lentiviral Packaging Mix. A complete media exchange was performed 24 hours post-transfection. Viral supernatants were harvested 72 hours after and centrifuged at 400*g* for 5 minutes to pellet residual virus-producing cells. The supernatant was then carefully transferred and aliquoted into assay plates, which were stored at -80 °C until use.

#### Lentiviral transduction

To generate transgenic 55yo cells, the 55yo pooled fibroblast population was transduced with lentiviral vectors encoding the ORF sequence of interest for each gene tested. On the day of transduction, fibroblasts were plated into the appropriate plate format in DMEM/F12 media supplemented with 15% Fetal Bovine Serum, 1% Penicillin/Streptomycin and 5*µg/ml* Polybrene. Viral supernatant was added to the plate. The plate was then left in the BSC for 10 minutes to allow for initial attachment of cells, and then spinfected at 400*g* for 90 minutes at 32°C. The cells were the transferred to the incubator overnight.

For scRNA assays, 1.52 *×* 10^5^ cells were transduced in 24-well format. For CellPaint assays, 672 cells were seeded in 384 well format. To ensure a pure population for scRNA assays, the transduced fibroblasts were selected under 2*µg/ml* blasticidin for 7 days. On Day 16 post transduction, fibroblasts were harvested and put through the ScalePlex fixation.

#### Single-cell RNA sequencing

Single-cell RNA sequencing was performed using the ScalePlex Single Cell RNA Sequencing Kit v1.1 (ScaleBio), following the manufacturer’s instructions. Cells were subjected to ScalePlex fixation and sample hashing. A total of 384 hashed samples were generated from four primary samples, each comprising 96 hashed conditions. All samples were thawed, counted, and normalized to enable equivalent cell loading across conditions.

Following a concentration step, cell suspensions were adjusted to a final density of 3, 500*cells/µL* for loading. Based on cell counts, approximately 17,500 cells were loaded per well into the reverse transcription (RT) plate for each sample. After RT, samples were pooled and distributed into a 384-well ligation plate. Ligation and subsequent pooling were performed according to protocol. The pooled libraries were then distributed into four final distribution plates, with 1,600 cells allocated per well. Samples were processed through the remaining steps of the ScalePlex workflow up to the Index PCR stage. Finally, a subset of PCR-amplified pooled libraries from each of the four distribution plates was taken forward for targeted enrichment.

Samples were sequenced on a Nexseq2000 with read lengths of 34 for read 1, 10 for index 1, 10 for index 2, and 76 for read 2. There was no additional PhiX spike in for this sequencing.

#### scRNA data processing and quality control metrics

After deconvolving the single-cell identities and associated perturbagens, we implemented a fairly standard program of data transformation and quality control. Our single-cell analysis was performed in the ScanPy library. Our processing and quality control procedure followed several sequential steps: removal of cells events thought to be empty or doublets (or higher-order multi-cell capture events); a calculation of the percentage of mitochondrial, ribosomal, and MALAT1 transcripts per-cell, with the intent to discard anomalous cells; a restriction of the library of genetic elements being quantified; normalization of the per-cell counts to counts-per-million (CPM); prediction of cell phase (though no regressing-out of the predicted phase or related genes was performed, nor was the data transformed in any way based on imputed cell-cycle stage); removal of mis-labeled control cells; finally, discarding from consideration all library members with low cell coverage.

Doublet and empty cell prediction and removal occurred at multiple stages. The ScalePlex procedure has an in-built scrubbing step for non-singlet events. Additionally, we used Scanpy’s **filter cells** command to remove cells with fewer than 1,000 overall counts.

The relative per-cell abundances of mitochondrial, ribosomal, and MALAT1 transcripts was calculated via the **calculate qc metrics** command with the alphanumeric “MT-” prefix associated with mitochondrial genes, the prefixes “RPS” and “RPL” associated with ribosomal genes, at “MALAT1” associated with MALAT1. Because the ScalePlex system is a combinatorial indexing system, which does not involve immediate lysing of the cell, the overall abundances of these transcripts (particularly mitochondrial and MALAT1) was quite low compared to other single-cell sequencing technologies. Ultimately, we saw no outlier cells in terms of these abundances, and did not discard any cells based on their relative QC-type transcipt abundances. Rather (or, perhaps more accurately put, “jointly”), as we argue in the paper, these are informative biomarkers.

The above procedures fixed the set of cells we considered viable to be included in analysis. We next restricted the set of genetic elements under consideration. Using the GrCH38 genome build and associated annotations, we discarded all transcripts associated with genetic elements that did not fall into two annotated categories: “protein coding” and “processed pseudogene”. The combined total of these categories summed to 18,609 distinct genetic elements whose expression was considered for analysis. After this restriction, the remaining transcripts associated with each cell were normalized to counts-per-million (CPM) using Scanpy’s **normalize total** command. The distinction between the usage of CPM normalization here, whereas we had argued TPM normalization was necessary for our implementation of CCM, amounts to the distinction between physics-based tools and statistical tools. Our primary usage of this scRNA data is to establish differential expression of genes under various perturbations, as well as machine-learning tools to cluster and classify single cells, neither of which are particularly sensitive to whether expression counts are meant to represent the exact number of underlying RNA molecules in the cell or a statistical proxy.

After these procedures fixed the experimental data matrix – consisting of rows (cells) by columns (genes) and their numerical entries – we took a few additional measures to try to avoid certain confounding effects that might exist in the data for downstream analysis. First, we scored each cell according to its imputed position in the cell cycle. This was done using the Scanpy **score genes cell cycle**, using a commonly-used panel of 96 genes strongly associated with the **S** and **G2/M** phases.^76^ Notably, we did not *regress out* these features from the data; that is to say, the numerical entries in the experimental data matrix did not change as a result of our cell scoring. The purpose of the scoring was to ensure that a cell’s current phase of cycle was not a confounder for any analyses (as established, for instance, via Kendall correlation between imputed cell cycle state and any other categorical imputed quantity such as young-like, or a cluster ID). We never observed such correlations in our analysis.

Lastly, two steps of particular salience to our intended analyses were performed. Firstly, the raw sequencing data was examined and aligned to a reference file containing non-native segments of our delivery plasmid. This was done in order to determine if any cells which, after deconvolution, fell into the 25yo/55yo control population, were expressing our plasmid. Such cells were therefore either mis-annotated, or had errantly received plasmid during handling. We found that 1,641 such cells (out of 62,885 55yo control cells and 54,642 25yo control cells; therefore 1.3% of control cells) and removed them from the experimental data matrix.

Finally, out of 283 perturbations which were present in the experimental data matrix at this stage, 17 ^4^ were judged to have especially low cell coverage (fewer than 100 cells; see Fig. S3). These cells and perturbations were removed from the experimental data matrix, leaving 266 unique cell conditions (which includes 25yo and 55yo untreated controls) distributed among 62,885 55yo controls, 54,642 25yo controls, and 532,319 perturbed cells.

#### Differential expression analysis

The structure of our differential expression analysis proceeded as follows: for our 264 uniquely-perturbed populations of 55yo cells, we performed an analysis via comparison to our untreated 55yo cell population to determine which genes were statistically significantly different in expression as a consequence of the genetic perturbation.

Differential expression analysis was performed using the Scanpy **rank genes groups** command with default hyperparameters and options. This means that the per-gene statistics associated with each of the 264 tests emerges from a Welch *t*-test on “pseudobulked” control and perturbed populations. This comports with the general understanding that pseudobulked methods provide the best measure of differential expression, even when single-cell information is present.^49^ In the absence of technical replicates, other pseudobulk methods such as EdgeR^77^ and DeSEQ^78^ are either inapplicable to this problem or are disadvantageous in the absence of replicates. Multiple-hypothesis correction (Benjamini-Hochberg) was implemented on all emerging *p*-values.

#### Removing the footprint of viral transduction and blasticidin selection

The act of lentivirus transduction – independent of genetic payload – is known to have a substantive impact on cell phenotype.^79,80^ The same is true for drug selection via an inclusion in the plasmid conferring resistance to blasticidin, followed by treatment with blasticidin after viral transduction.^81,82^ These effects present significant confounders for us, as any “overlapping” phenotypic and transcriptomic changes induced by our individual library perturbations which can be attributed to the pure act of transduction and subsequent drug selection will skew our analytical results.

The extent of this background effect can be seen in the number of differentially-expressed genes in the cell population with GFP transduced. We found that 10,536 genes (56.7% of all genes analyzed) were differentially expressed in this population compared to the 55yo baseline. Since GFP is generally treated as an inert control, we took this large set of genes to be essentially equivalent to those which are perturbed by lentiviral transduction, irrespective of payload.

Our method of treatment for this confounding effect was simply to omit this panel of 10,536 genes from downstream analyses, rather than remove them from the experimental data matrix (which would, among other consequences, render cells unnormalized). This was implemented in two ways: for a comparison of the topology of networks inferred by CCM and revealed by differential expression analysis (next section), we simply omitted all GFP-induced differentially-expressed genes from any entry in the contingency table for the Fisher Exact test; in our derivation of a transcriptomic measure of rejuvenation (subsequent section), we did not permit any trained model to use as feature any gene differentially expressed in GFP-perturbed samples.

#### CCM network accuracy and precision analysis

We measured the similarity between the CCM-inferred network and the differential expression results (which we construe as a measure of CCM’s accuracy in the manuscript) in two fashions. The first, a global measure, is simply the overall accuracy of the network – for our preferred cutoff on the CCM strength at full library, *ρ\** ⩾ 0.5677, this accuracy is the number of CCM-inferred edges (i.e., set relationships of causer and causee) which were confirmed by differential expression analysis divided by the total number of CCM-inferred edges which *could possibly have been validated* by the differential expression (causer-causee pairs where both genes exist in the initial discovery dataset as well as our scRNA experimental dataset For this precise CCM implementation, the accuracy would be 53.8% with all experimentally-measured genes included, and 30% with the GFP differential expression profile removed. Repeating the same analysis for two comparanda methods, DeepSEM^52^ and the method of partial correlations^50,51^ gave accuracies of: DeepSEM: 42.4% with all genes, 22% with the GFP differential expression profile removed; Partial Correlations: 48% with all genes, 22.3% with the GFP differential expression profile removed.

For both cases (with and without the genes contributing to a “viral expression profile”), we computed similar accuracies for a null hypothesis that we took to represent a reasonable facsimile of random chance. This null was constructed as follows: for the 263 relevant library genes in our experiment, we recorded the out-degree distribution in the inferred CCM network. A single realization of the null hypothesis consisted of first drawing a focal gene *g* from the 263 library genes, then drawing an out-degree *d_g_* from this discrete distribution uniformly-at-random and without replacement, then selecting *d_g_* unique genes as edge recipients from this focal gene from the set of all genes *G_tot_* = *G_CCM_ ∩G_expt_*, the genes which co-occur in both the pilot dataset and our experimental dataset. This null is intended to represent a randomly-constructed network which “topologically resembles” the CCM-inferred network, but with causer-causee relationships shuffled. This choice of degree distribution and set from which to pick recipient genes (nodes) gives high precision for the random null because of the very high number of differentially-expressed genes that we generally saw in our perturbed cells, even when the “viral profile” is removed from consideration. For Figure 2, 100,000 realizations of the null hypothesis were computed to give the precision distributions shown.

A broader implementation of this global measure of CCM performance is given by the precision-recall curve at variable cutoffs *ρ\**, rather than the cutoff *ρ* >* 0.5677 which we had selected as the primary implementation in the rest of the manuscript. This curve is plotted in Fig. S4. In both the general case and the case used in the manuscript, with the background differential expression profile for viral transduction removed, the curves have virtually identical shapes. In both cases, the optimal precision is obtained at a much more lenient value of *ρ\** than we utilized in our main-use network. An increase in precision of 5%-7% would come from keeping, instead of the top 5% of edges (thresholded at *ρ\** = 0.5677), rather the top 65% of edges (thresholded at *ρ\** = 0.052. This threshold is somewhat interesting. Recall that these quantiles are calculated only over pairs of edges that satisfy the *convergent* property of CCM – that the *p*-value of the Spearman correlation of CCM strengths *ρ*(*L*) and library sizes *L* be *p_Spear_ <* 0.05. Seeing optimal performance at a low value of *ρ\** – just greater than 0, the minimum possible value for the “cross-mapping” property of CCM to be valid – suggests that any strict thresholding above the base definition of the method leads to a decrease in performance.

Finally, a critical measure of method success is also given by a local (i.e., per-gene) measure of accuracy. It is reasonable to expect that the set of interactions imputed to one gene will cryptically be much higher in accuracy than the set imputed to another – though we have attempted to quantify the inductive bias of the method throughout our work, there are certain to be dynamic and biological associations of genes for which this method is more applicable than for other genes. For our local test, we treated the comparison between CCM-inferred interactions and differential expression relationships as a contingency table, mirroring the way pathway enrichments are typically calculated. The contingency table for any focal gene consists of four categories: (A), 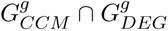, the set of recipient genes *G* for a focal gene *g* which were both inferred by CCM and confirmed via differential expression; (B), 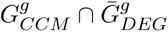, the set of inferred relationships that were <u>not</u> confirmed via differential expression; (C), 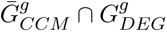, the set of relationships found by differential expression which were not predicted to exist by CCM; and (D), 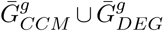, the set of all validateable genes which were not associated with the focal gene *g* by either CCM or differential expression. The statistical fidelity of the CCM network was then measured on a per-gene basis as the odds ratio and *p*-value of the Fisher Exact test associated with the gene’s contingency table.

#### scRNA binary classification and derivation of the YouthScore

Our method for measuring rejuvenation contrasts somewhat with other recent methods in the literature, such as “aging clocks”, which take in biological data from a cell or individual and return a continuous-valued prediction of the chronological (or “biological”) age of the sample. Our logic for this deviation was based on both mathematical and biological considerations. The biological considerations emerge directly from our experimental structure, which in turn were informed by the practicality of cell availability from existing biobanks – it is far easier to curate two disparate aging populations, such as our mixed 3-donor pools of 25yo and 55yo cells, than to curate sufficient differently-aged individuals (whilst ensuring experimental compatibility between different donor cells, such as comparable passage number, cell health, etc). These control populations could now reasonably be expected to reflect the inter-donor variability in the two differently-aged pools, the intra-donor single-cell heterogeneity, and any cell behavior induced by our library perturbations which did not cleanly align with either of these unperturbed categories.

We also believe that the appropriate mathematical framework for this problem will now be less susceptible to noise (and overfitting) and more interpretable than aging clocks. The decreased susceptibility to noise comes through the binary nature of the problem; a well-trained model will classify most of the 55yo test cells correctly as 55yo, capturing the intra-donor variability, and naturally leading to lower generalization error; a continuous-valued regression model, on the other hand, can be more susceptible to generalization error by producing training accuracy which is “close enough”, but still contains strong donor-specific model features (i.e.: donor A cells can consistently be called 52yo, donor B 55yo, and donor C 58yo; the average error will be small enough for the model to be considered “good”, but there is a latent association of cells with individual donors rather than the quantitative age). The high-codimension nature of aging clocks (thousands of genes regressed onto 1 continuous variable, the imputed age) also permits a very wide set of disparate models which are “just as good” as one another in terms of performance; that is, their Rashomon basin^83^ is likely quite large. We believe this hampers the interpretability of the features *driving* the accuracy. Lastly, we believe the type of binary classification framework we employ is more suitable for our library experiments, where perturbed 55yo cells can reasonably be expected to look *neither* like endogenous 25yo or 55yo cells, but rather like a novel synthetic phenotype; this comes through our implementation of abstention (see below).

As an initial exploration into what type of binary classification framework to employ, we trained all of the applicable scikit-learn binary classifiers with base hyperparameters and settings on our experimental data matrix and examined the train-test accuracies on unperturbed 25yo and 55yo cells. Few methods seemed to train well on this data, with Gaussian, Bayesian, and SVM-type models achieving accuracies on class-balanced samples which were between 50% and 60%. The highest performance was observed with decision-tree type classifiers, particularly random forest and AdaBoost.^84^ Ultimately, we determined to orient our framework around AdaBoost, due to its ease of adaptibility to aspects of our problem and its inherent regularizing activity.^54^ We valued the latter property as both improving interpretability (because the age-determining features are fewer in absolute number, and can also be associated with the evolution of example (cell) weights in each round of boosting) and, when employed in a bagged ensemble of models (see below), hedging against Rashomon-type effects by ultimately sampling important features from a varied subset of examples (put differently: if a particular subset of cells – and no others – and their associated model show a strong aging association with gene X, then, with sufficient subsampling and bagging, the importance of X will be proportional to the size of the distinguished cell population).

The two adaptations we made to basic implementations of AdaBoost (i.e., that with per-example exponential weights, reweighted at every step of boosting^54,84^), were the additions of bagging and per-bagged model abstention. Bagging (or “bootstrap aggregating”), involves sub-sampling the set of available examples in a way that balances the number of training examples from each class (and associated hold-out test examples), which then serve as the training space for a single model. When repeated, this procedure generates a “bag” of models, which can vary in their determining features and training-example-predictions, even on overlapping training examples. This has obvious advantages in terms of the “representational” qualities of the bag of models in comparison to any single model, which might for instance be susceptible to be “overpowered” by a small set of outlier examples. The subsampling is also necessary to achieve class balancing in imbalanced datasets such as ours (since, as would naively be expected, our 25yo fibroblasts divided at a faster rate than our 55yo fibroblasts). We viewed class balancing as an advantage for maximizing our per-example margins, as well as enhancing the positive effects of voter abstention. In our work, individual AdaBoost models were run for 300 rounds of boosting each, and the ensemble consisted of 10 such models, each with equal voting weights.

We believe voter abstention both harmonizes well with our bagging approach and addresses the issues that come from our end goal – using the trained classifier ensemble to evaluate 55yo cells which have been perturbed by overexpression of library members. The primary concern here is that the perturbed cells are unlikely to come from the same probability distribution over features as endogenous, “natural” 55yo and 25yo cells; on the contrary, they could (and often did) seem *un*natural in comparison to these populations. If this degree of “unnaturalness” was taken to an extreme, it would vitiate our entire approach; efforts to maximize our training accuracy and interpret our models would be meaningless, as the generalization error would be decoupled completely from the training error.^84^ The meaninglessness of the exercise could in principle be difficult to spot; without abstention, voters are constrained to return per-example class probabilities for only 25yo and 55yo, with no ability to communicate the alien nature of the example cells under consideration. Abstention also hedges against outlier cells arising either from intra- or inter-donor heterogeneity. Even unusual endogenous cells will often be abstained upon by one or more models (in fact, examining the number of training cells abstained upon as a function of the abstention margin is how we fix the latter hyperparameter).

We implemented abstention via existing schemes.^85^ At a per-voter level, abstention occurs when the class probability *p_c_*of an example is *|p_c_ −* 0.5*|* ⩽ *A*, where A is the abstention margin. The factor of 0.5 arises from the class balancing activity of our bagging. In our work, to determine A, we tracked the fraction of training cells upon (averaging over all 10 voting models), *f_C_* = *ι_|pC −_*_0.5_*_|_*_⩽_*_A_/N*, as a function of *A*, which was evaluated over a grid [0.005,0.0075,0.01,0.0125,0.015,0.0175,0.02,0.0225,0.025]. We also tracked the total margin *M* over non-abstained examples, 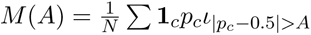, where **1***_c_* indicates whether or not the called class (25yo or 55yo) is correct for the example in question. Both quantities are plotted in Fig. S6**(a)**. A principled way of choosing the abstention rate might be to choose the “elbow” according to one or both curves and fix the value of *A* accordingly. However, we also admit a certain bias towards abstaining on fewer training cells when possible, simply to try to keep the available training data as large and broad as possible. In the end, examining both curves, we fixed *A* = 0.01 for our scRNA work.

After the bagged ensemble model was trained and the abstention margin fixed, we began the process of evaluating each individual perturbed 55yo cell, *i*. Each of the 10 voting models *j*, for each of this cell, emits an associated class probability 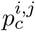, as well as a vote *v_ij_* – a selection from “25yo” 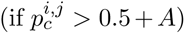, “55yo” 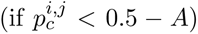, or “I don’t know” 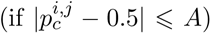. Denoting these outcomes (and their associated frequencies among the 10-voter ensemble, as a shorthand) as *v*_25_*_yo_*, *v*_55_*_yo_*, and *v_A_*, the ensemble as a whole emits at least two relevant quantities; the majority vote *v_M_* = max([*v*_25_*_yo_, v*_55_*_yo_, v_A_*]), and the averaged class probability 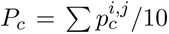. Ultimately, we decided to use the majority vote as the source of our measure for rejuvenation. We defined this “YouthScore” *Y* as *Y* = *ι_v__M_* =*v*_25*yo*_/*N_Ā_*, where *N_Ā_* denotes the set of all cells where the majority vote *v_M_* was not abstention, *v_A_*. Said differently, the “YouthScore” is the frequency of *false 25yo calls* by the entire classifier ensemble on cells which are, in truth, perturbed 55yo cells.

#### CellPaint experiments

Our CellPaint^86^ assay was performed at the Broad Institute. Fibroblasts were transduced following the protocol described above in the scRNA experimental methods section. 5 days after transduction, the cells are stained with phenovue641 mitochondrial stain for 30 minutes, then washed with Hanks Balanced Salt solution, and fixed for 20 minutes by paraformaldehyde.

Phenoview dyes are reconstituted in 1x PhenoVue Diluent A. The following are combined as a multistain cocktail per 384 well plate:: 75*µL* 20% Triton X-100, 18.75*µL* 568 (phalloidin), 15*µL* Hoechst 33342, 150*µL* 555 WGA, 18*µL* 512 (SYTO14) nucleic acid stain, and 37.5*µL* 488 concanavlin A. The cells permeabilize and stain with this multistain coocktail for 30 minutes at room temperature. The cells are washed again with Hanks Balanced Salt solution, before imaging on the Perkin Elmer Opera Phenix instrument. Imaging is done in non-confocal mode with the 20x water objective, with 9 fields taken per well.

#### CellPaint data processing

The structure of our single-cell morphological CellPaint data is very similar to scRNA data – the experimental data can be represented by a matrix, will rows consisting of cells and columns consisting of morphological features which are extracted from raw, stained images using the CellProfiler software.

In contrast to scRNA data, there is little in the way of “canonical” data quality control and normalization procedures for CellPaint. Concerns such as empty/damaged cells, doublets, and higher-order multiplets generally do not exist. If certain measured features are indicative of cell health and integrity at time of assay (such as mitochondrial and ribosomal count densities in scRNA), these are not well-known or well-documented in the literature.

As a consequence, we included all cells in the experimental data matrix. However, we did perform feature engineering on the columns of the matrix. Unlike transcriptomic data, where any multicollinearity (strong correlations between columns) is an interesting result reflecting biological connections between the expressions of two or more genes, multicollinearity in our CellPaint data is likely to emerge simply as a consequence of the way the CellProfiler package is programmed and utilized. A cursory look at the CellProfiler manual reveals the classes of features that can be extracted from images, as well as the methods available to calculate these features. Many CellProfiler pipelines, such as ours, will employ all methods listed which are compatible with computing a given feature. Naturally, these methods can often give extremely similar results. For instance, the module *MeasureImageIntensity* – which is meant simply to quantify the amount of dye in each channel in the cytoplasm, nucleus, and whole cell, respectively – consists of the methods/measurements *TotalIntensity, Mean Intensity, MedianIntensity, StdIntensity, MADIntensity, MinIntensity, MaxIntensity, LowerQuartileIntensity, and UpperQuartileIntensity*. It is easy to visualize how some of these measures can be extremely well-correlated with one another. Other modules, such as those that measure spatial correlation between dye channels, will often use similarly defined methods for measuring spatial correlation, which can be numerically near-equivalent in many circumstances.

This amount of multicollinearity is undesirable for our supervised learning applications, because it significantly increases the odds of overfitting. It will also vitiate the margin-maximizing properties of AdaBoost, since it increases the ability of the algorithm to select multicollinear features as focal features, which will effectively decrease its ability to exploit the full feature set when the maximum number of rounds of boosting is bounded (which is always the case in practical applications). Useful and novel information, such as the features with the highest Gini importance in determining youthful and aged cells, can similarly be drowned out with collinear features.

To combat this effect, we pared the feature set as follows: for each of the 6,812 features in our initial data matrix, we calculated the Pearson correlation with every other feature. We decided on a cutoff of *ρ_P ear_* ⩾ 0.9 above which two or more features would be considered excessively collinear (i.e., redundant). To ensure that our final data matrix contained no combinations of features that were excessively collinear, we iteratively removed these features from the data matrix. For instance, call the first feature examined *A*; if features *E* and *Z* are collinear with *A*, they are removed from the data matrix at this stage (and we annotate that the feature *A*, when used in analysis, is standing also as statistical proxy for *E* and *Z*). Proceeding thus through every feature renders a data matrix with no excessive collinearity. This matrix will have a slight dependence on the order in which features are examined; shuffling the features before the procedure will lead to different choices of the proxy members for sets of collinear features. Our justification for this is that the cutoff *ρ_P ear_* ⩾ 0.9 is rather high; the higher the cutoff, the less overall numerical difference can be observed between one realization of the data matrix achieved via this procedure and another one with the order of the features shuffled.

We performed an additional step in feature engineering, related to the arrayed nature of the assay. Because our chief analysis on the CellPaint data will be a repetition of our binary classification (“YouthScore”) performed on our scRNA dataset, it is critical that our pooled 55yo and 25yo control cells show no bias in morphological featurization based on their location in a particular well or on a particular plate. The CellPaint assay could be subject to different sources of technical noise, both at a per-well level (with edge wells being subject to increased evaporation of medium as well as potential optical biasing events) and per-plate level (variations in total dye and other wet-handling variables). Morphological features that reflect this variation would be then less reliable for classifying young and aged cells. To combat this, we measured the coefficient of variation of every feature (after temporary *z*-score normalization) across both the 55yo control cell population and the 25yo control cell population. By examining the distribution of the coefficients of variation (Fig. S7), we determined a cutoff of *CV* ⩽ 0.3 to include.

Our final data matrix consisted of 139,532 55yo control cells; 41,162 25yo control cells; and 685,914 cells spread over 278 distinct library perturbations; for each cell, 4,421 morphological features which showed low variance among 55yo cells, low variance among 25yo cells, and no excessive collinearity with any other feature.

#### CellPaint binary classification and YouthScore

Our analysis of the CellPaint data was extremely similar to the binary classification and ensuing “YouthScore” measure of rejuvenation derived based on our transcriptomic data. To combat the class imbalance in our control cells (where we had roughly three 55yo cells for every one 25yo cell), we implemented bagging by training ten class-balanced AdaBoost models. Each model was run for 100 rounds of boosting; the test accuracies for the ten models were 85.2 *±* 0.3%. All models used between 50 and 60 distinct features, and the overall Gini importance of features pictured in Fig. 4 was the mean Gini importance over all ten trained models.

As in our transcriptomic work, we then quantified aging in each cell by presenting it to all ten trained models and recording the mean class probability (the consensus probability that the cell was a true 55yo cell, or was mis-identified as a 25yo cell). In a similar vein to our scRNA work, by examining the mean per-cell classifier margin and abstention fraction as we varied the abstention margin *A*, we determined that *A* = 0.0125 was an acceptable abstention margin for this work (Fig. S6 **(b)**). If the cell had a mean class probability (where the mean is measured over all models which did <u>not</u> abstain on the focal cell) that was higher for a 25yo cell than for a 55yo cell, it was deemed a false positive. The annotated “YouthScore” for each genetic perturbation is the fraction of false positives, i.e., the fraction of perturbed cells erroneously classified as 25yo despite being 55yo plus a transduced genetic perturbation.

#### *Post hoc* attempts to discover an improvement to the RankScore

We investigated several methodologies in an attempt to discover an alternative function on the non-experimentally-derived data (i.e., the GTEx raw data, its dynamic featurization via tsfresh, and the CCM-inferred network and network-derived featurizations) which would correlate better with our observed YouthScore(s). Here we describe attempts to fit the transcriptomic YouthScore; our attempts on the morphological YouthScore rendered similar results.

The methodologies used were all regression-based methodologies, attempting to regress the available features onto the measured YouthScore(s). They included: LASSO^71^ with 5-fold cross-validation (implemented via the scikit-learn package’s ‘LassoCV’ command with base hyperparameters); decision trees^87^ (implemented via the scikit-learn package’s ‘DecisionTreeRegressor’ command with base hyperparameters and a 70/30 train/test split); AdaBoost^84^ (here, implemented via the scikit-learn package’s ‘AdaBoost-Regressor’ command with base hyperparameters and a 70/30 train/test split; note that this contrasts with most of our AdaBoost implementations in the rest of the manuscript); linear SVM (implemented via the scikit-learn package’s ‘svm’ command with base hyperparameters and a 70/30 train/test split. Finally, we trained a graph convolutional network. This architecture (PyTorch Geometric) consisted of three graph convolutional blocks (all of width of 32), each followed by batch normalization, rectified linear unit (ReLU) activation, and dropout layers. The final layer consists of a 1-unit graph convolution without activation, for regression. Optimization used Adam (learning rate 10*^−^*^3^, weight decay rate 10*^−^*^3^) with mean squared error loss for 2,000 epochs.

Our network-derived features take two essential forms: node-based statistics, such as degree distributions, centralities, and other common measurements (all extracted from the two network topologies using the networkx); then, at a higher level, featurizations of the network using a Node2Vec^88^ embedding (64 components with a walk length of 30). Node2Vec was implemented via the nodevectors package.

Our dynamics-derived features included the features derived from the processed *quasi* -time series via tsfresh, the *quasi* -time series themselves, and the intrinsic featurization used in the base RankScore (the slope of the best-fit line to the *quasi* - time series, times (1 *− p_Spearman_*), where the *p*-value is derived from the Spearman correlation between the *quasi* - time series and the time axis).

All methodologies were applied to four distinct cases, permutations of two network topologies and two featurization schemes. The two topologies considered were the base CCM network used in the manuscript, that with CCM *ρ* >* 0.5677; the second topology was the same as the first, but restricted to the experimentally-tested “well-mapped” subnetwork that had a Fisher Odds ratio associated *p*-value of 0.2 or less. The first featurization included all available dynamic and network-derived features; the second featurization included only the “intrinsic” dynamic features used in the base RankScore and the Node2Vec features. For both featurization schemes, we considered both an unscaled feature set, and a *z*-scored feature set (implemented via the scikit-learn “StandardScaler” command).

We note that there is a very large imbalance between the size of the feature set and the amount of data being regressed upon, no matter whether the smallest or largest network topology and featurization scheme are employed. The YouthScore(s) are only known for *<* 300 genes, whereas the featurizations are available for thousands of genes. This had obvious consequences for the performance of the different methodologies tested; it was also a driving motivation for the methodologies we chose to use, which we view as spanning different tiers of model complexities (with LASSO at the lowest complexity and the graph convolutional network at the highest complexity). As expected, the more complex models struggled due to the dearth of training data. The graph convolutional network showed an erratic epoch-to-epoch objective function score, and in all cases failed to converge to a successful model.

The other methodologies, in some train-test configurations, landed on productive models. With the exception of LASSO (because of the difference in train/test splits; LASSO had a 5-fold cross-validation scheme, whereas all other methodologies employed a 70/30 single-fold train-test split), all methodologies were implemented 100 times, and the results averaged. These averages are plotted in Fig. S9. Most models intermittently found strong models, but in most train-test configurations failed to converge to a productive model (meaning one with an *R*^2^ ⩾ 0). LASSO, however, converged on a model which produced *R*^2^ *≈* 0.1 when applied to the smaller featurization on the “well-mapped” network. This is an incremental improvement on the base RankScore, which has *R*^2^ = 0.0841; however, it comes with *p <* 0.05, unlike the base RankScore. We believe it is clear that more training data would be necessary for any regression methodology and featurization approach to achieve better results.

## Acknowledgements

AHG was supported by grant DP5OD019851 from the Office of the Director at the National Institutes of Health. MSK, DM and DT were supported by an Wyss Institute Project award. We thank Alison Hill, Jeff Gerold, and Martin Nowak for helpful comments and suggestions.

## Contributions

MSK, DM, and DT designed the study. MSK, AHG and GS designed the CCM analysis. MSK and YF designed the AdaBoost-driven analyses. MSK implemented the CCM analysis and all downstream analysis. DM, DT and GC designed the experiments. DT, NC, BP and RF implemented the experiments. MSK, TW and WG analyzed the results of experiments. All authors wrote the manuscript.

## Data and code availability

Raw and processed scRNA data is hosted on Geo as GSE350289. A released codebase that replicates key methods and figures may be found on Github.

## Correspondence

All inquiries may be forwarded to MSK.

## Abbreviations

CCM: convergent cross-mapping
25yo: 25-year-old
55yo: 55-year-old
ER: endo-plasmic reticulum

### Appendix

#### Appendix A Code Availability

All relevant code will be made available upon publication.

#### Appendix B Supplementary Figures and Tables

**Figure S1.**
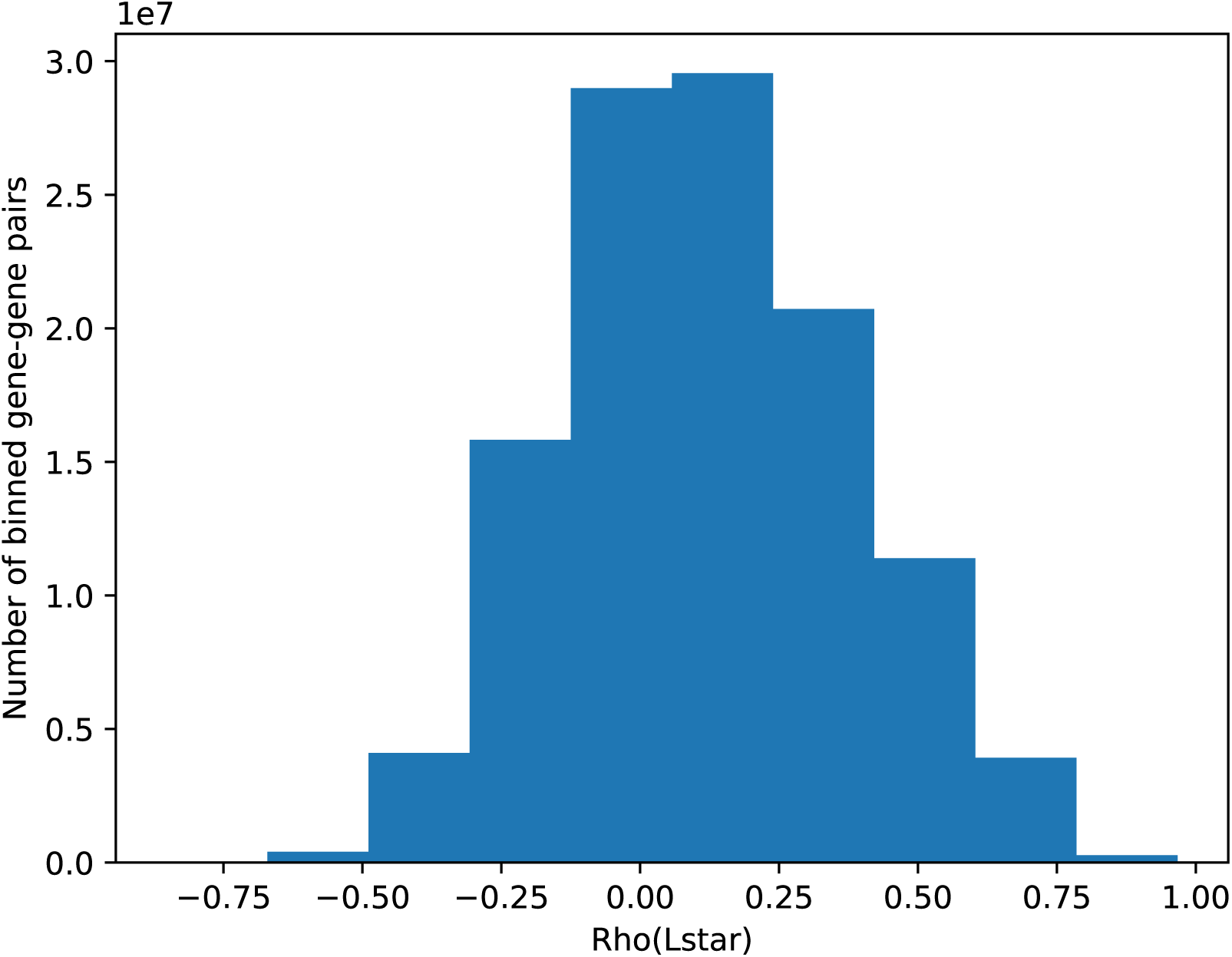
A histogram of the nearly 36 million gene-gene pairs examined by CCM. For each pair, the “CCM strength” – the maximum *ρ*(*L\**) (as a function of embedding dimension *E*) – is the value pictured. This distribution is very close to the normal distribution.

**Table S1.**
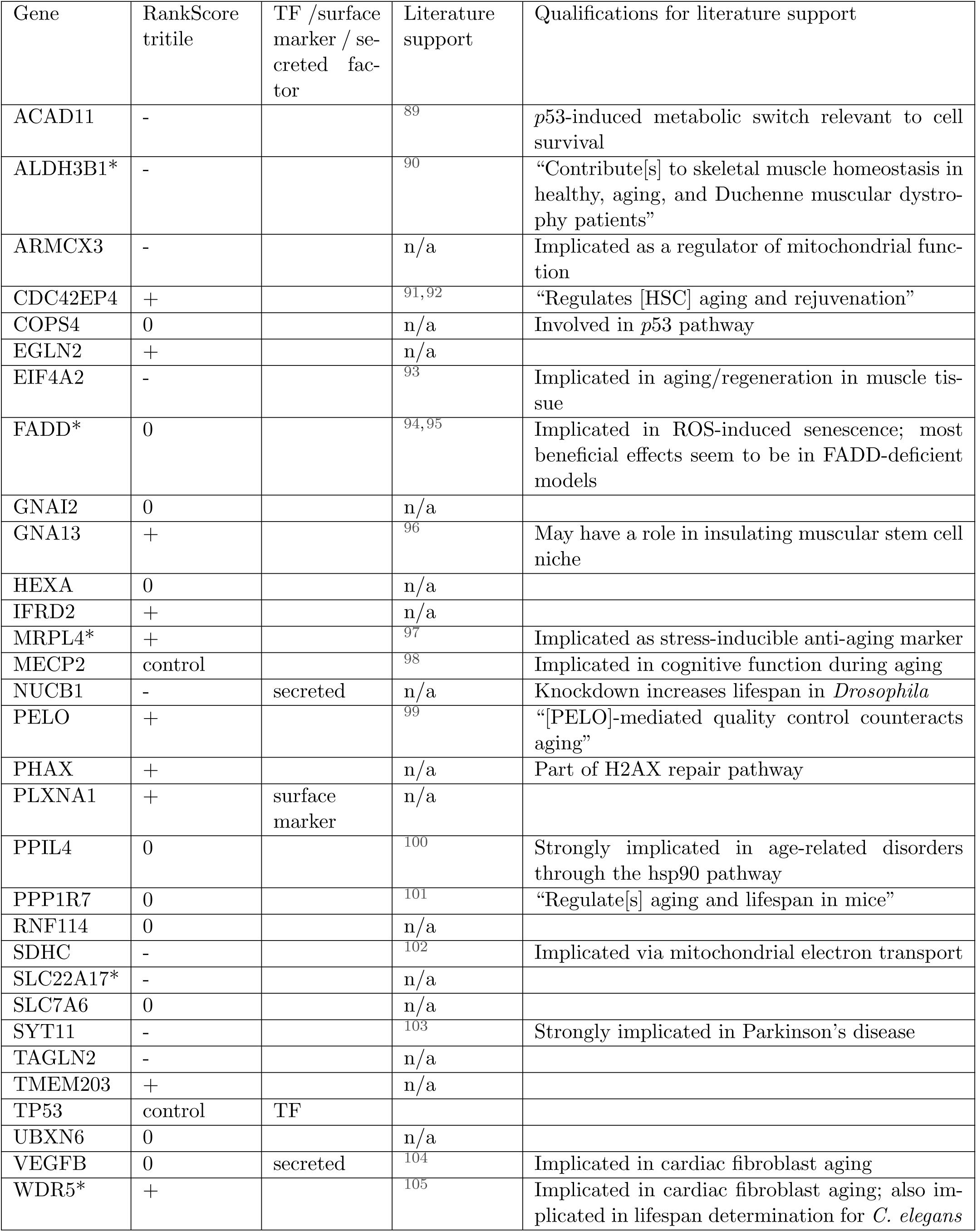
A short literature review of top-performing genes in this study. Genes included here showed rejuvenative effects in both our scRNA and CellPaint studies (excepting those with an asterisk * following their HUGO gene name; these fell in the top 5 by rejuvenative effect in only one of our assays).

**Figure S2.**
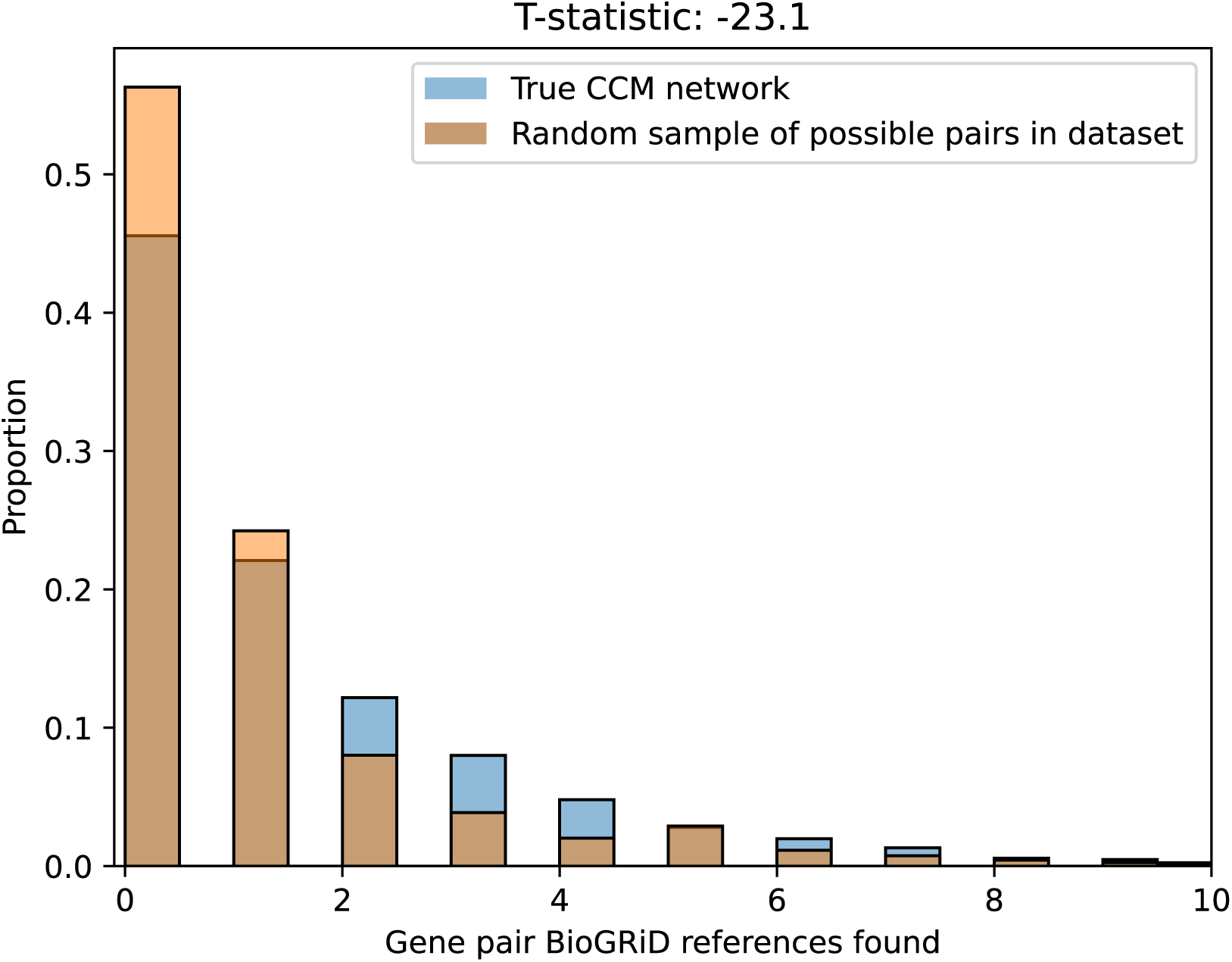
A comparison of BioGRiD citation distributions between our CCM network, consisting of roughly 1.8 million edges between roughly 6,000 genes, and a random drawing of the same size from BioGRiD – an alternative set of 1.8 million edges chosen uniformly-at-random from the *∼* 36 million possible edges amongst our genes. The random sample shows far lower support in the literature.

**Figure S3.**
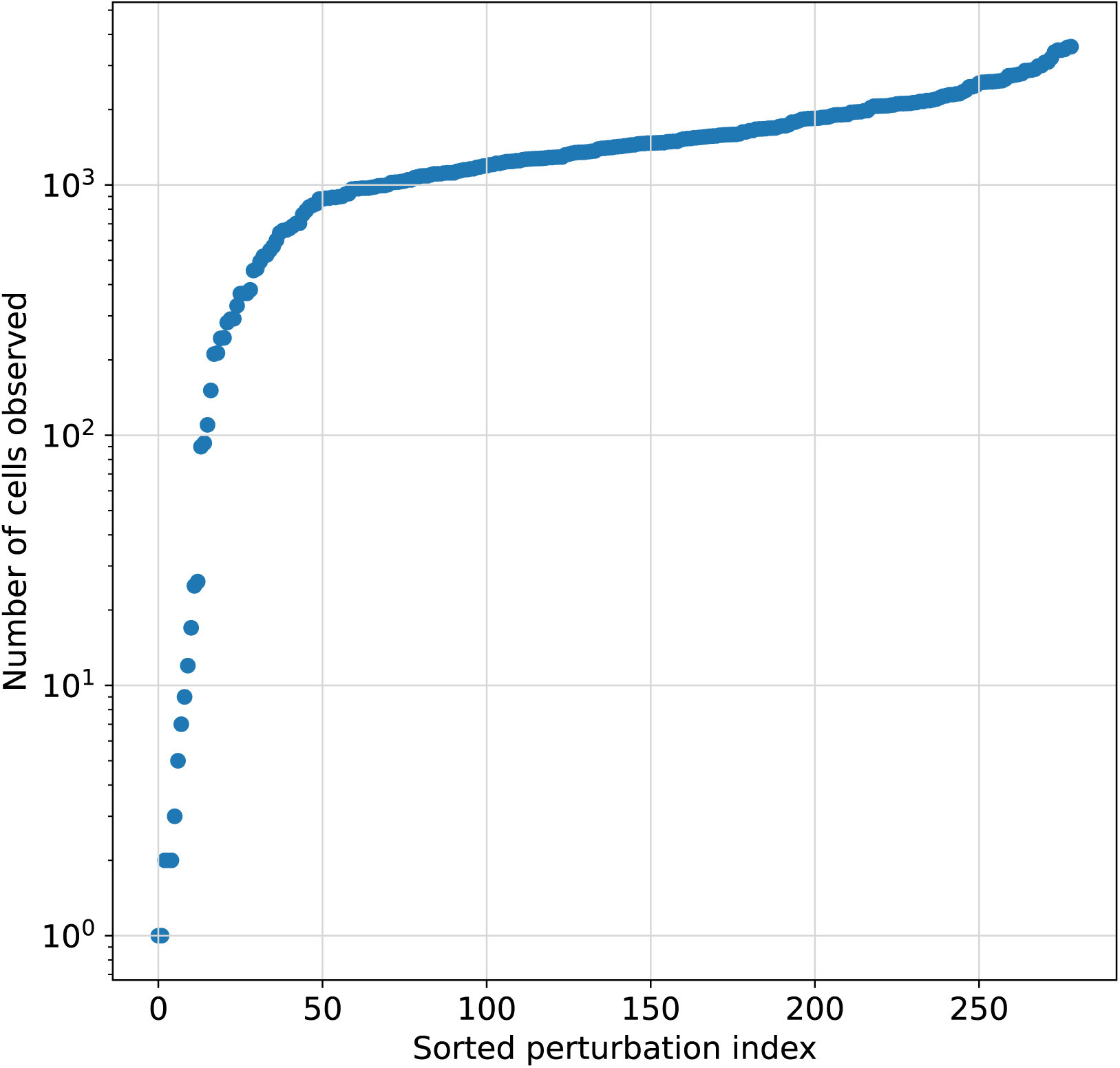
Cell counts per library perturbation after scRNA quality control measures. The 17 library members with cell counts less than 100 were omitted from further analysis due to low coverage.

**Figure S4.**
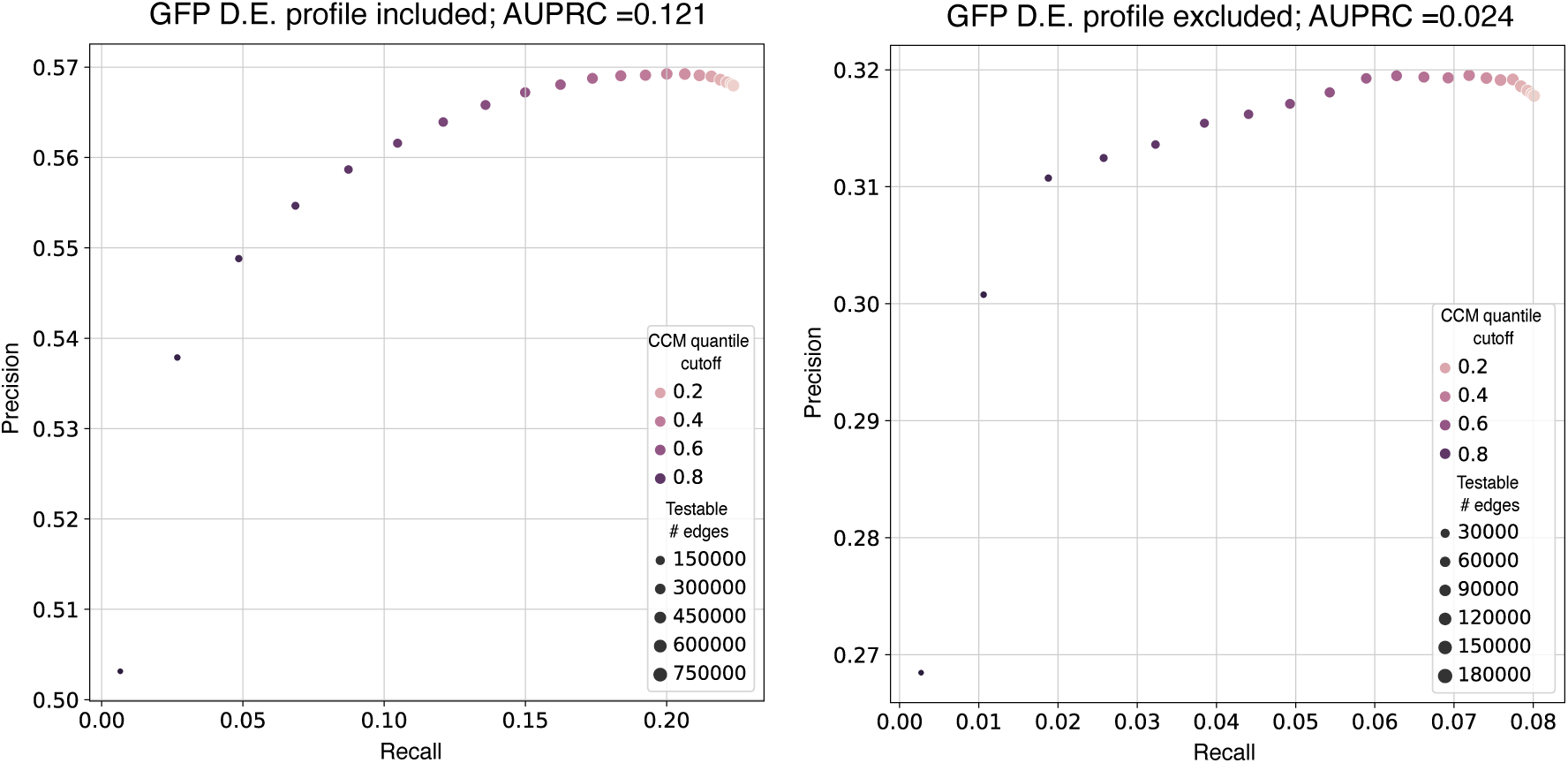
Precision-recall curves for the CCM network at varying cutoffs of *ρ_L∗_* (*X, Y*), using the differential expression profiles as the ground truth, with the observed GFP differential expression profile (the viral transduction profile) either included or excluded.

**Figure S5.**
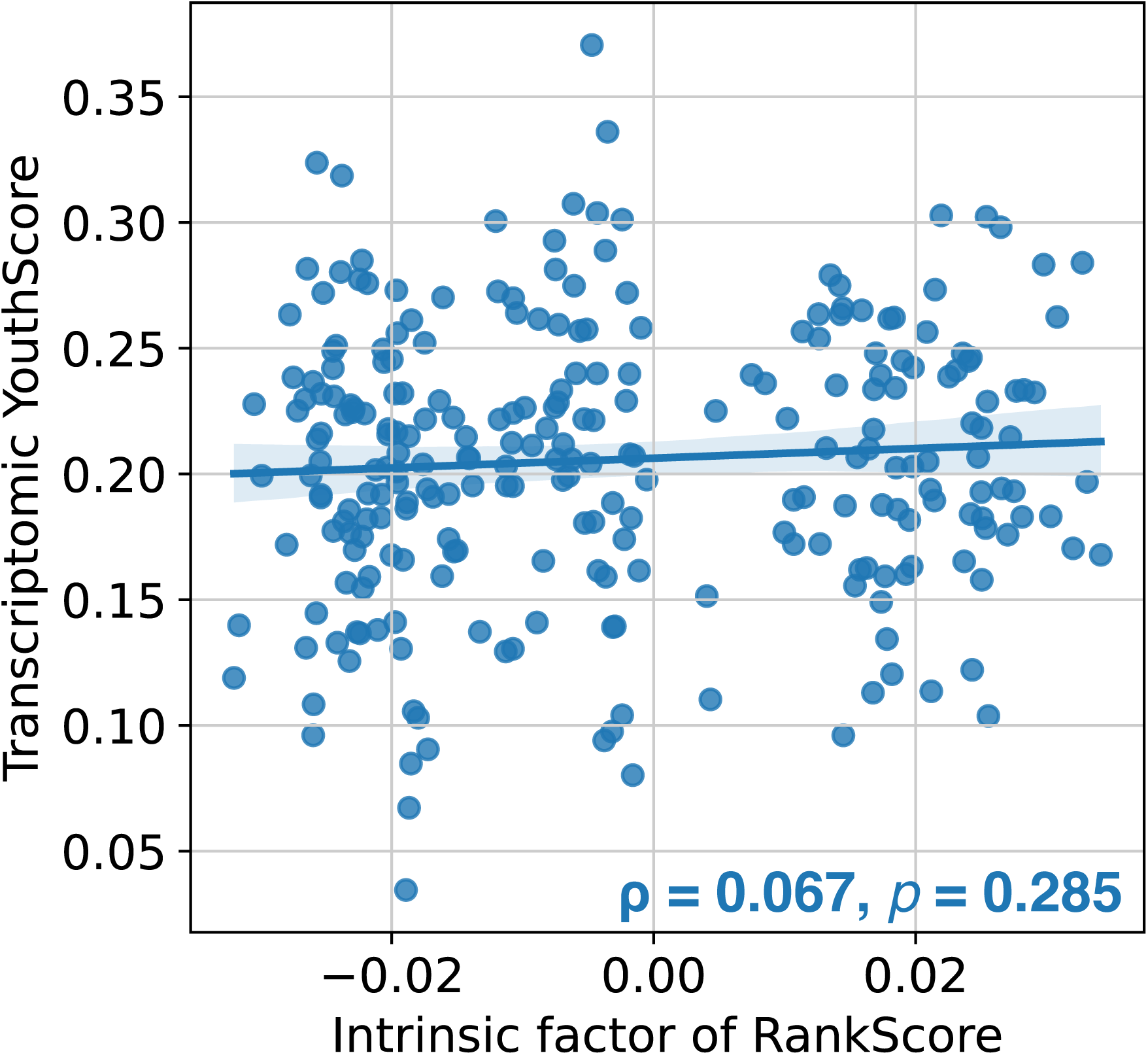
A regression plot of the intrinsic per-gene factor of the RankScore – whose sign is given by the slope of the best-fit line to the *quasi-* time series data – versus the transcriptomic YouthScore.

**Figure S6.**
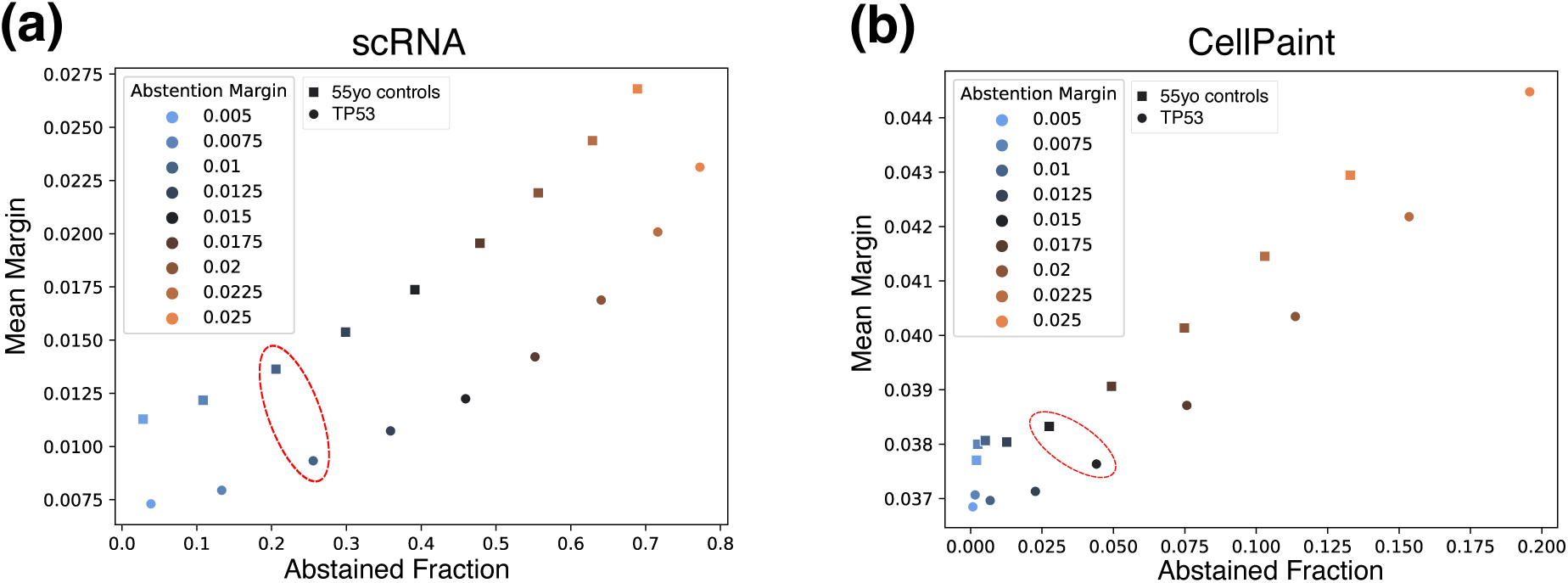
**(a):** The mean per-cell margin of the ensemble scRNA classifier, versus the fraction of cells for which the majority vote was abstention. These were calculated on an evenly-spaced grid of abstention margins *A* (colors), and we used their behavior on the 55yo controls (square markers) to determine the preferred value of *A* (circled, red). To show how these quantities might behave for library perturbations, we show identical curves for TP53 (circles). **(b):** The same as **(a)**, but for our CellPaint work. Both quantities are generally improved compared to scRNA, with higher per-cell margins and relatedly lower rates of overall abstention.

**Figure S7.**
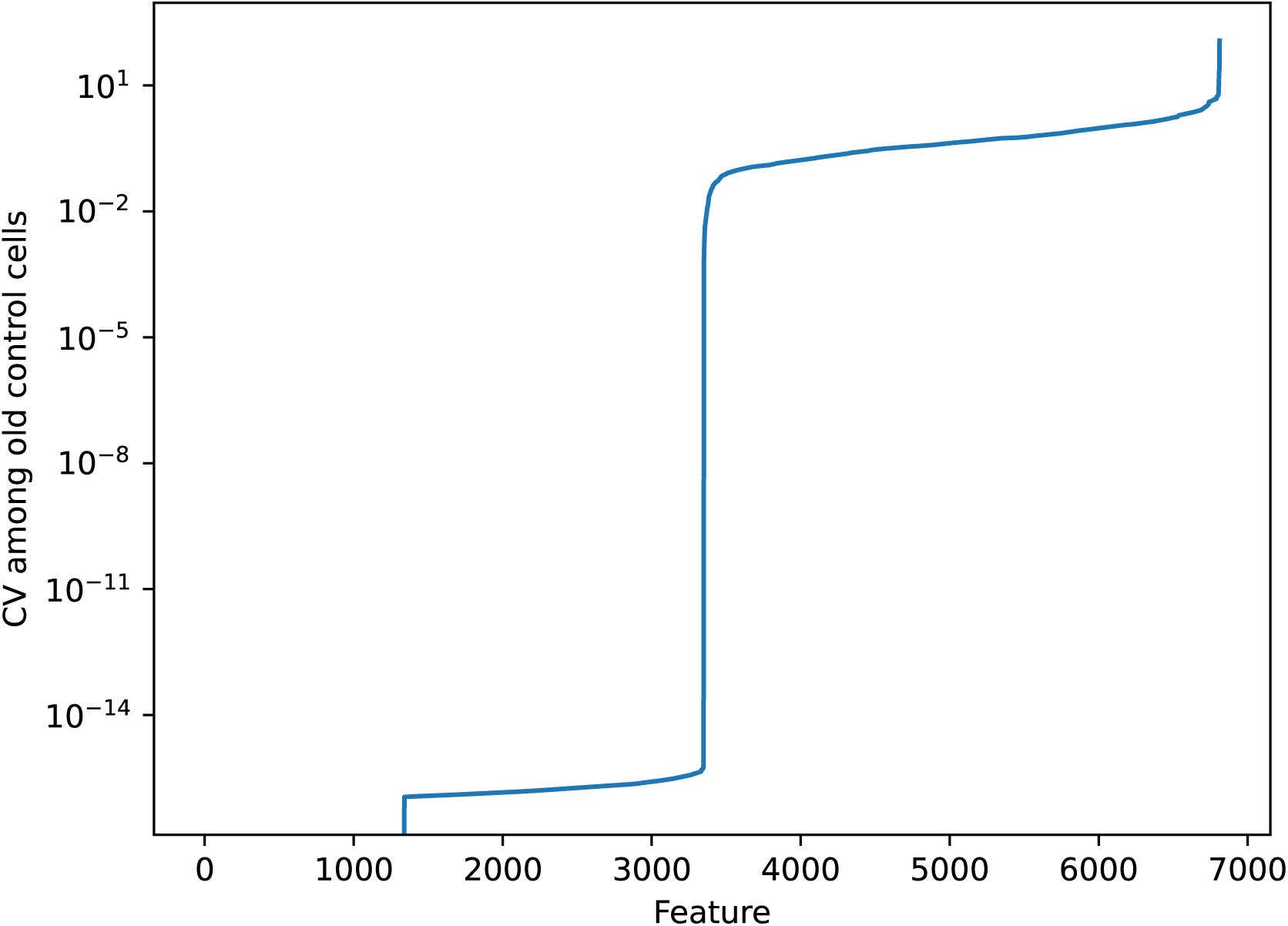
A sorted line plot of the CellPaint feature coefficients-of-variation (CV) across old control cells. For the CellPaint analysis, features with *CV >* 0.3 (the elbow in this plot, as well as the identical plot for young control cells) were omitted from analysis.

**Figure S8.**
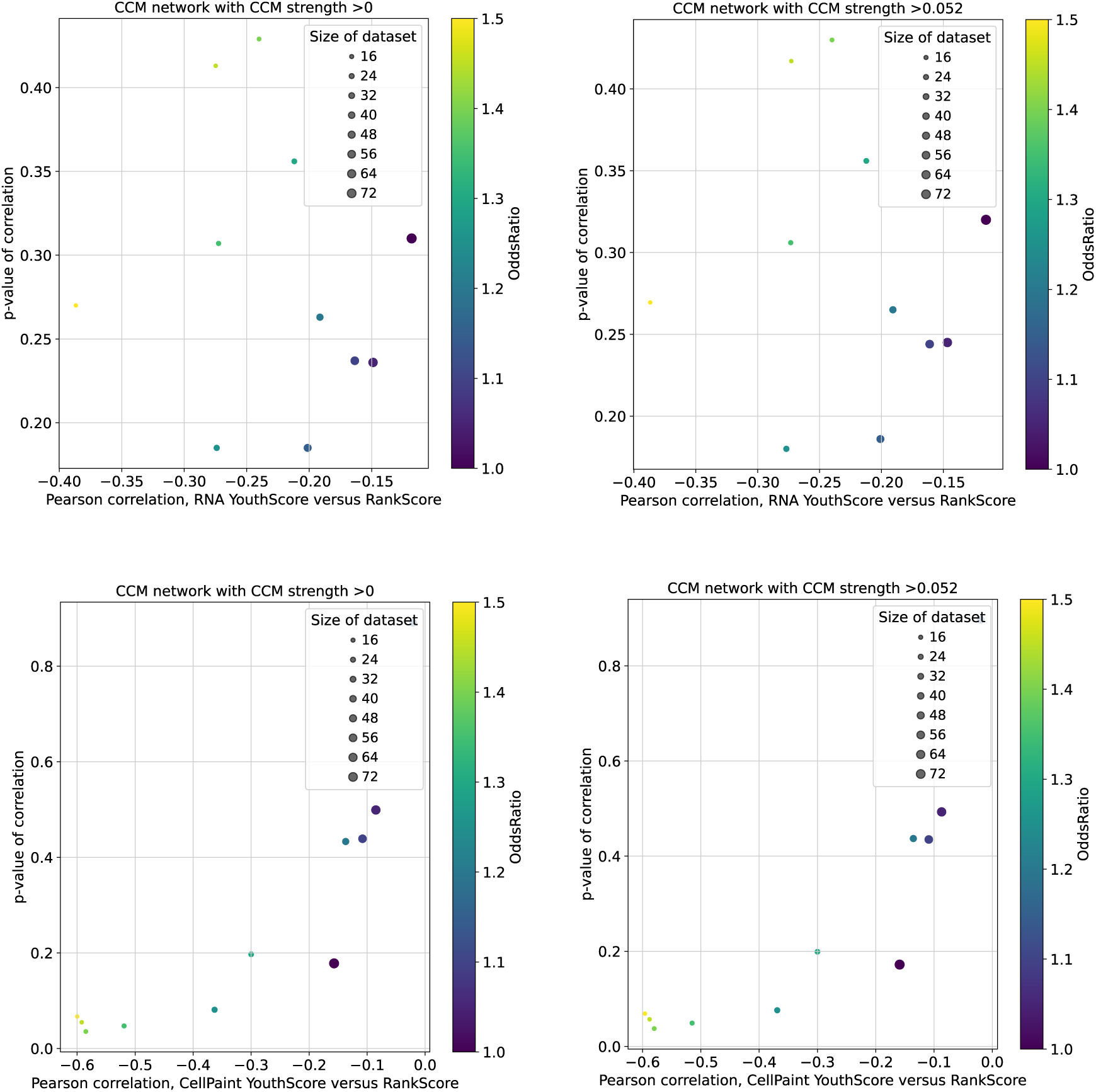
An examination of how RankScores, derived according to Equation 1 but with larger and more permissive CCM topologies, correlate with the observed YouthScores drawn from both scRNA and CellPaint. The two larger networks under consideration are those which are derived from **(LHS)**: any gene-gene pair which shows the convergent property of CCM (Spearman-associated *p*-values of library size and CCM strength at statistical significance), and show any end-library cross-mapping score *ρ* >* 0; **(RHS)**: the optimally-precise network with *ρ* >* 0.052 (see Fig. S4). Pearson correlations between these RankScores and the YouthScores for scRNA (top) and CellPaint(bottom) are given on *x* and the corresponding *p*-values on *y*; in every case, we consider an ablation of “well-mapped” gene neighborhoods based on the Fisher odds ratio, and mark the odds ratio cutoff and resulting library size accordingly. Whereas the scRNA-derived YouthScore does not improve in correlation with the RankScore, the CellPaint-derived one does, showing extremely strong and significant correlation with the broader RankScores, especially in the most well-mapped gene neighborhoods.

**Figure S9.**
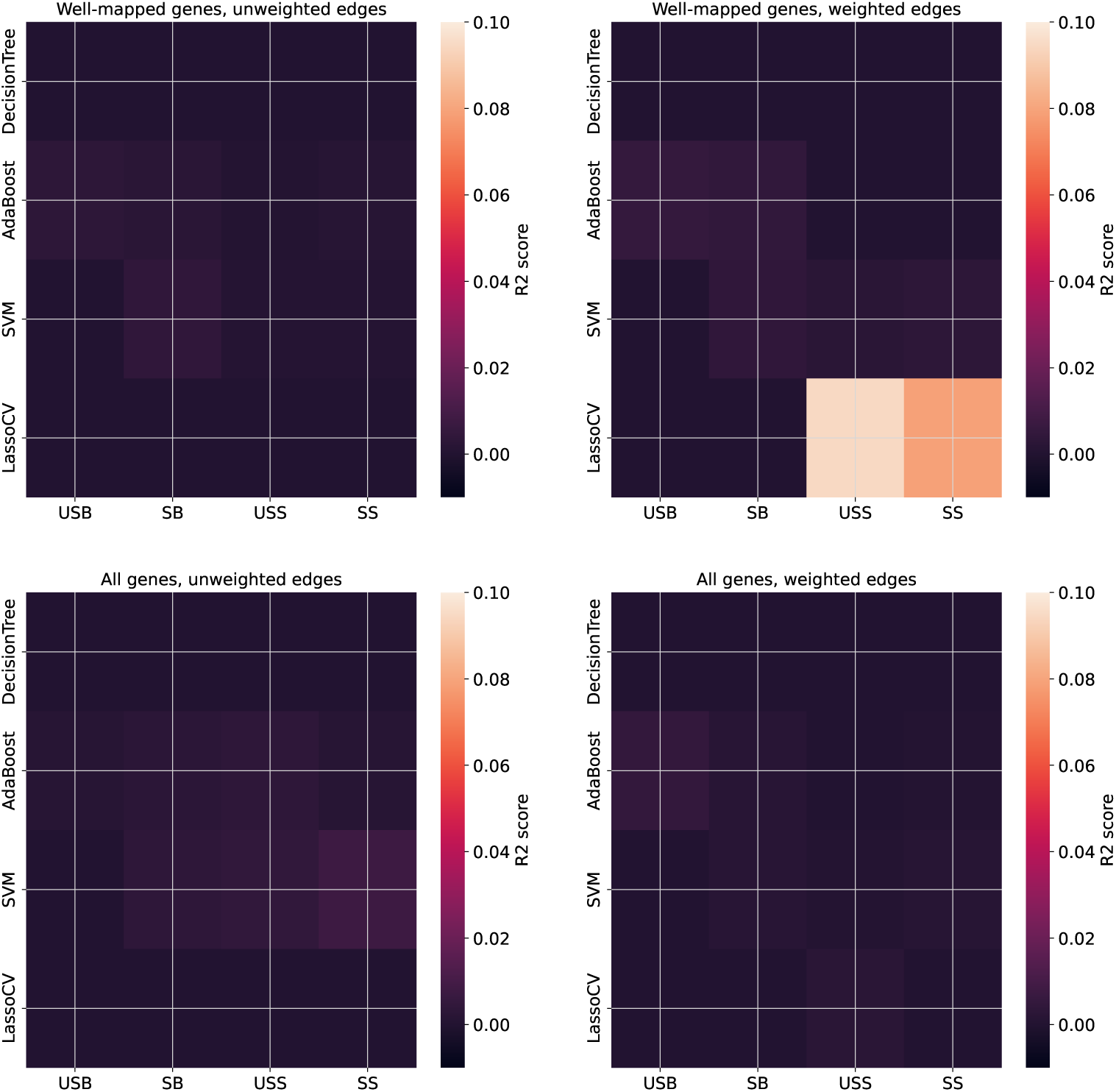
An initial exploration of alternative RankScores found using additional featurization and augmentation of the CCM network. We explored four network topologies, all with the *ρ* >* 0.5677 cutoff implemented throughout the main text – (A) only well-mapped gene neighborhoods according to our scRNA analysis; (B) the same, but with edges weighted by CCM end-library strength *ρ\**, as opposed to our unweighted edges; (C) the network precisely as implemented in the main text, (D) the full main-text network but with *ρ\**-weighted edges. For each topology, we explored four classes of models (decision trees, Adaboost, linear SVM, and LASSO) as applied to four novel featurizations arising from a binary yes-no on feature *z*-score scaling and arising from either a ’big’ or a ’small’ feature set. ’Small’ feature sets included **Node2Vec** featurizations of the underlying topology, as well as the best-fit slope of the *quasi-* time series. ’Big’ feature sets additionally included **tsfresh** featurization and the entire *quasi-* time series. The featurizations are labeled “USB”=’unscaled, big’; “SB”=’scaled, big’; “USS”=’unscaled, small’; “SS”=’scaled, small’.

**Figure S10.**
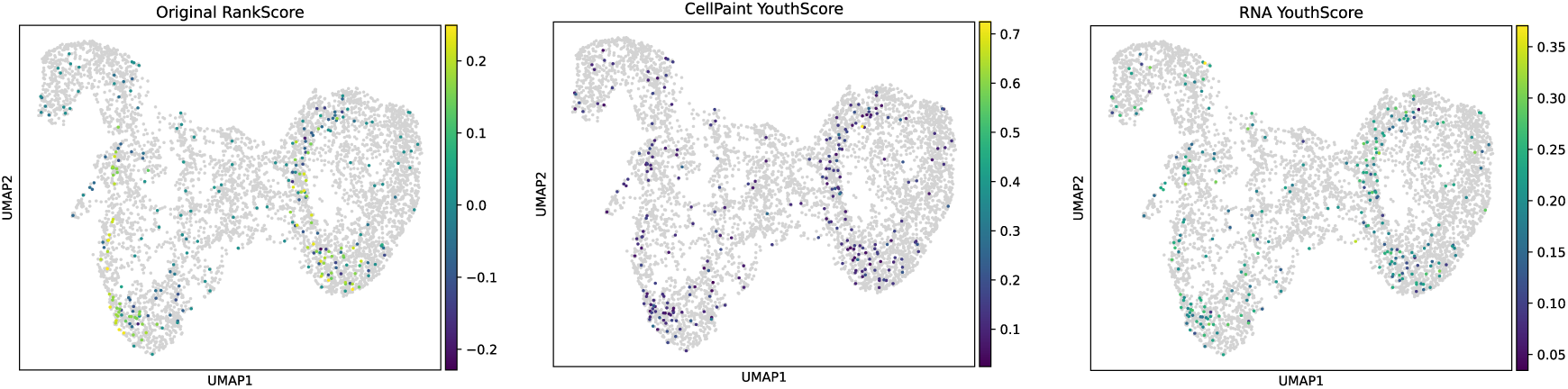
Alternative colorings of UMAPs in Fig. 5 with key quantities used throughout the manuscript.

## Footnotes

1 CREB3, ZGPAT, ATF6B, SMAD5, REPIN1, NCOA3, CEBPG, MXD4, ZKSCAN1, CXXC1, EBF1, ZNF768, CAMTA2

2 PLXNA1, VAMP5, SLC3A2, CAPNS1, SLC29A1, LMAN2, MXRA8, KITLG, CD68, SIDT2, ITGA4, S1PR3, LRRC32

3 PVR, NUCB1, VEGFB, FN1, LY96, RNASE4, LGALS1

4 USO1, H3F3B, HNRNPK, DNAJB12, YARS, RAB23, RER1, EFTUD2, DDX49, TEX261, GABARAP, SUPV3L1, NFIX, FADD, CDC42EP3, MARK3

